# Multi-Cohort Blood Transcriptomic Meta-Analysis of Active Pulmonary Tuberculosis: Reproducibility of a 50-Gene Interferon Signature and Benchmarking of a 4-Gene Classifier

**DOI:** 10.64898/2026.11.19.767834

**Authors:** Usama Hassan, Farah Tahir, Armeela Bibi, Warda Jamal, Tooba Kanwal

## Abstract

**Background:** Most blood transcriptomic signatures for tuberculosis (TB) come from a single population or a single platform. Few have been built by meta-analysis and benchmarked against published signatures.

**Methods:** We meta-analysed three UK blood transcriptomic datasets (two Illumina microarray, one RNA-seq; 247 samples: 87 active pulmonary TB (PTB), 160 healthy controls (HC)). GSE107991 reanalyses the Berry test set, and two discovery datasets overlap Sweeney’s discovery data, so the datasets are not fully independent. Genes were retained if they replicated in all three studies (FDR < 0.05, same direction) and passed random-effects meta-analysis (|logFC| > 1.5, FDR < 0.01). Performance was estimated in GSE28623 (The Gambia, n = 83) and GSE56153 (Indonesia, n = 36); direction was checked in GSE34608 (Germany, n = 26) and GSE81746 (India, n = 6). A LASSO and random-forest classifier was compared with the Berry 393-gene and Sweeney 3-gene signatures. Immune composition was assessed with MCPcounter and CIBERSORTx.

**Results:** We identified a 50-gene core signature dominated by interferon-stimulated genes. All 50 were upregulated and reached FDR < 0.05 in every dataset. ssGSEA gave an AUC of 0.830 (95% CI 0.743–0.917) in GSE28623 and 0.941 (0.868–1.000) in GSE56153. Direction agreement was 95.7% in Germany and 95.8% in India. PTB versus latent TB infection gave 0.848 (0.756–0.940). LASSO selected four genes (*BATF2*, *ANKRD22*, *GBP5* and *EPSTI1*). The frozen classifier gave an external AUC of 0.864 (0.785–0.944). The Sweeney 3-gene score, re-tested in one of its validation cohorts, reached 0.912 and exceeded the 50-gene score (47/50 genes; Holm *p* = 0.0234). Berry 393-gene scoring gave 0.910 with directions reconstructed from GSE19439 (Holm *p* = 0.0071 versus the 50-gene score). The signed score is a reconstruction, not Berry’s published score. MCPcounter showed myeloid expansion and lymphoid suppression; CIBERSORTx recovered the former but not the broader lymphoid suppression.

**Conclusions:** The 50-gene signature was consistent across three overlapping UK discovery datasets. External evaluation was partial, comprising AUCs for 47 of the 50 genes in two cohorts and direction concordance in two smaller cohorts. This is a benchmarking study, not a claim of superiority: the Sweeney 3-gene and signed Berry scores were higher, and the 4-gene classifier did not differ significantly from Sweeney. The classifier warrants prospective validation. The Berry and Sweeney comparisons are not independent.

## 1. Introduction

Tuberculosis (TB) remains the leading cause of death from a single infectious agent worldwide, with an estimated 10.7 million new cases and 1.23 million deaths in 2024 [1]. Diagnosis of active pulmonary TB (PTB) still rests mainly on sputum: smear microscopy, culture and nucleic acid amplification tests. Each needs a respiratory specimen, laboratory infrastructure and trained staff [1]. In high-burden, resource-limited settings, where most cases occur, these requirements are hard to meet. A blood-based test that avoids sputum would help most in smear-negative patients and where microbiology facilities are out of reach.

Blood transcriptomics is among the most studied non-sputum approaches to TB biomarkers. Berry et al. (2010) described a 393-transcript signature in whole blood, interferon-inducible and neutrophil-driven, that separated active TB from healthy controls and from latent TB infection (LTBI) [2]. That work established the interferon-stimulated gene (ISG) module as the dominant feature of the systemic host response to *Mycobacterium tuberculosis*. Sweeney et al. (2016) used multicohort analysis to derive a 3-gene diagnostic signature (*GBP5*, *DUSP3*, *KLF2*) that discriminated active TB from controls across several datasets [3]. Zak et al. (2016) took a different route, identifying a 16-gene signature that predicted progression from LTBI to active disease [4]. Singhania et al. (2018) organised the response into co-expressed modules, which captured phenotypic variation across the disease spectrum [5].

Most signatures have been derived and validated within a single cohort by internal cross-validation. Multicohort derivation is established [3], but prospective testing in geographically or ethnically distinct populations remains uncommon. Cross-platform integration is also uncommon. Microarray and RNA-seq TB datasets now coexist, yet they are rarely combined in one meta-analysis. A third gap is mechanistic: statistically derived signatures are seldom checked against the cellular changes that accompany active TB, particularly the myeloid expansion and lymphocyte suppression seen in peripheral blood.

We meta-analysed three UK datasets (two microarray, one RNA-seq; 247 samples after quality control). One of them, GSE107991, is a reanalysis of the London test set used to validate the Berry 393-gene signature, so these are three discovery datasets, not three independent cohorts. Genes were selected in two stages, first by per-study replication and then by random-effects meta-analysis, which left a 50-gene core signature. We tested that signature without further tuning in four external cohorts from The Gambia, Germany, India and Indonesia; two were large enough for AUC estimation and two supported only a direction check. We also trained a 4-gene classifier, compared it with published TB signatures, and used deconvolution to ask which cell types carry the signal. Discovery and evaluation were kept separate, and Section 4.7 discusses the limits of that separation.

## 2. Materials and Methods

### 2.1 Study Design and Datasets

We performed a multi-cohort blood transcriptomic meta-analysis to identify a consensus gene expression signature for active pulmonary tuberculosis (PTB) versus healthy controls (HC). Three discovery datasets were analysed: GSE83456 (Illumina HumanHT-12 v4 BeadChip; United Kingdom) [6], the Bloom dataset (GSE42826 + GSE42830, sub-series of GSE42834; Illumina HumanHT-12 v4 BeadChip; United Kingdom) [7], and GSE107991 (Illumina HiSeq RNA-seq; United Kingdom; Singhania et al. 2018) [5]. The discovery datasets spanned two platforms (Illumina microarray and RNA-seq) and all came from UK cohorts.

GSE107991 is an RNA-seq reanalysis of the 54 samples in GSE19444, the London test set in which Berry et al. validated their 393-transcript signature [2]. All 54 GSE107991 samples are annotated in the Gene Expression Omnibus (GEO) as reanalyses of GSE19444, and their GEO source names begin Berry_London_Test_set_. Its 28 post-QC samples are therefore not independent of that validation cohort (Table S18). The studies share no subject-level identifier that would permit cross-referencing, so patient overlap among the UK datasets cannot be excluded; we therefore describe them as three discovery datasets rather than three independent cohorts. This overlap does not change the within-study replication criterion, but it limits the independence of the head-to-head comparisons discussed in Section 2.11.

External evaluation used four held-out cohorts. None informed gene selection, thresholding, or model tuning. Two cohorts supported AUC estimation: GSE28623 (Agilent GPL4133; The Gambia, West Africa; Maertzdorf et al. 2011) [8] and GSE56153 (Illumina HumanRef-8 v3, GPL6883; Indonesia; Ottenhoff et al. 2012) [11]. Two smaller cohorts supported direction-concordance analysis only: GSE34608 (Agilent GPL6480; Germany; Maertzdorf et al. 2012) [9] and GSE81746 (Agilent SurePrint G3 Human GE v2 8×60K, GPL17077; India; Sambarey et al. 2017) [10]. GSE28623 and GSE34608 come from the same investigator group.

### 2.2 Cohort Definitions and Sample Composition

Latent tuberculosis infection (LTBI) samples were excluded from all discovery and primary validation analyses to focus on the PTB-versus-HC diagnostic contrast. Specifically, 21 LTBI samples were removed from GSE107991 before quality control, and 25 LTBI samples were removed from GSE28623 before the primary validation. LTBI samples in GSE28623 were subsequently re-incorporated for a dedicated PTB-versus-LTBI sub-analysis (Section 2.10).

Post-quality-control composition of the discovery and validation cohorts is given in Table 1.

**TABLE 1.** Discovery and Validation Cohort Characteristics.

| Dataset / Cohort | Platform | Population / Geography | Initial / Deposited Samples | Exclusions and Quality Control | Analysed PTB (n) | Analysed HC (n) | Total analysed (n) | Study role |
|---|---|---|---|---|---|---|---|---|
| **Discovery Cohorts** |  |  |  |  |  |  |  |  |
| **GSE83456** | Illumina HumanHT-12 v4 BeadChip (GPL10558) | United Kingdom (London) | 106 (45 PTB, 61 HC) | 3 outliers removed (>3 SD from centroid: 2 PTB, 1 HC) | 43 | 60 | 103 | Discovery (microarray) |
| **Bloom** (GSE42826 + GSE42830) | Illumina HumanHT-12 v4 BeadChip (GPL10558) | United Kingdom (London) | 117 (27 PTB, 90 HC) | 1 HC outlier removed (>3 SD from centroid) | 27 | 89 | 116 | Discovery (microarray) |
| **GSE107991** | Illumina HiSeq 2500 (RNA-seq) | United Kingdom (London) | 54 (21 ATB, 21 LTBI, 12 HC) | 21 LTBI excluded (design); 5 author outliers removed (4 ATB, 1 HC) | 17 | 11 | 28 | Discovery (RNA-seq); reanalysis of Berry test set GSE19444 |
| *Discovery Subtotal* |  |  | *277 deposited* | *30 excluded (21 LTBI, 9 outliers)* | *87* | *160* | *247* | *Pooled Discovery Pool* |
| **Validation Cohorts** |  |  |  |  |  |  |  |  |
| **GSE28623** | Agilent Whole Human Genome Oligo 4x44K (GPL4133) | The Gambia (West Africa) | 108 (46 PTB, 37 HC, 25 LTBI) | 25 LTBI excluded from primary validation; 0 outliers removed | 46 | 37 | 83 | Primary external validation (ssGSEA AUC) |
| **GSE34608** | Agilent Whole Human Genome 4x44K v2 (GPL6480) | Germany (Europe) | 44 (8 PTB, 18 HC, 18 sarcoidosis) | 18 sarcoidosis disease controls excluded; 0 outliers removed | 8 | 18 | 26 | External validation (direction concordance) |
| **GSE81746** | Agilent SurePrint G3 Human GE v2 8x60K (GPL17077) | India (South Asia) | 8 (4 PTB, 1 pooled TB, 1 LTBI, 2 HC) | 1 pooled TB (non-independent) and 1 LTBI (design) excluded; 0 outliers removed | 4 | 2 | 6 | External validation (direction concordance) |
| **GSE56153** | Illumina HumanRef-8 v3.0 Expression BeadChip (GPL6883) | Indonesia (Southeast Asia) | 71 (18 active PTB, 18 HC, 15 treatment, 20 recovery) | 15 mid-treatment & 20 post-recovery samples excluded; 0 outliers removed | 18 | 18 | 36 | External validation (ssGSEA AUC) |
| *Validation Subtotal* |  |  | *231 deposited* | *80 excluded (26 LTBI, 18 sarcoidosis, 35 treatment/recovery, 1 pooled)* | *76* | *75* | *151* | *Multi-Continent External Evaluation* |
| **Total Overall** |  |  | **508 deposited** | **110 excluded** | **163** | **235** | **398** | **Full Study Dataset Universe** |

**Notes**
1. **PTB:** Active pulmonary tuberculosis; **HC:** Healthy controls; **LTBI:** Latent tuberculosis infection.
2. **Discovery Design Exclusion:** LTBI samples in GSE107991 (n = 21) were excluded to focus on the PTB versus HC diagnostic contrast.
3. **GSE107991 Provenance:** GSE107991 is an RNA-seq reanalysis of the 54 samples in GSE19444, the London test set in which Berry et al. validated their 393-gene signature. All 54 samples are annotated in GEO as reanalyses of GSE19444. It is therefore not independent of that cohort.
4. **Primary Validation Cohort (GSE28623):** Primary validation used the 83-sample cohort (46 PTB versus 37 HC). The 25 excluded LTBI samples were evaluated in a separate sensitivity analysis of PTB-versus-LTBI discrimination.
5. **Validation Outlier Policy:** No outlier filtering was applied to any validation cohort; all deposited samples meeting the study group definitions were retained.
6. **GSE56153 Clinical Staging:** 15 samples during anti-TB chemotherapy and 20 post-treatment recovery samples were excluded before normalisation as post-baseline entities.

For GSE107991, the full GEO deposit contained 54 samples (21 ATB, 21 LTBI, 12 HC); 21 LTBI were excluded wholesale and 5 author-flagged outliers (4 ATB + 1 HC) were removed before quality control, yielding 28 samples. For GSE81746, 1 pooled TB sample (GSM2175022) was excluded because pooled RNA from multiple patients violates biological-replicate independence, and 1 LTBI sample (GSM2175023) was excluded consistent with the LTBI exclusion policy, yielding 6 samples. For GSE56153, 15 mid-course treatment and 20 post-treatment recovery samples were excluded before normalisation as biologically distinct categories, yielding 36 baseline samples. HIV status was not available in the GEO metadata for GSE56153. GSE28623, GSE34608 and GSE56153 were all used previously to validate the Sweeney et al. 2016 3-gene signature [3].

### 2.3 Quality Control and Preprocessing

#### Outlier detection

For the two microarray discovery datasets (GSE83456 and Bloom), outliers were identified as samples falling more than 3 standard deviations from the disease-group centroid in principal component analysis (PCA) space (PC1 + PC2). Three outliers were removed from GSE83456 (2 PTB + 1 HC) and one HC outlier was removed from Bloom. For GSE107991, 5 samples pre-flagged by the original authors via the GEO series matrix (outlier:ch1 column) had been removed prior to analysis; PCA on the remaining 28 samples detected 0 additional outliers. Quality control plots (PCA, density, boxplot, sample-correlation heatmap) were generated for each dataset before and after normalisation.

#### Bloom sub-series batch assessment

The Bloom dataset is a merge of two GEO sub-series (GSE42826 and GSE42830). Pre-normalisation PCA coloured by sub-series revealed no sub-series-driven clustering; PC2 (4.6% variance) separated disease status (PTB versus HC), whereas PC1 (50.4%) reflected overall variance and did not track batch origin. The two sub-series were therefore merged and treated as a single batch for normalisation. No ComBat or removeBatchEffect correction was applied.

#### Normalisation

| **Dataset** | **Strategy** | **Notes** |
|---|---|---|
| GSE83456 | Series matrix used as-is | GeneSpring-processed deposit |
| Bloom | Series matrix used as-is | Same platform/pipeline as GSE83456 |
| GSE107991 | DESeq2 internal size-factor normalisation | Raw integer counts confirmed |

The GSE83456 GEO deposit was GeneSpring-processed rather than raw, so limma::neqc() was not applied.

#### Validation dataset preprocessing

For GSE28623 and GSE34608, raw Agilent data were background-corrected using limma::backgroundCorrect(method = “normexp”) and normalised with limma::normalizeBetweenArrays(method = “quantile”) [12]. Probe-to-symbol collapse was performed using max-mean probe selection with platform-specific annotation (GPL4133 for GSE28623; GPL6480 for GSE34608), yielding 19,749 gene symbols for GSE28623 and 19,575 for GSE34608. Sample exclusions (LTBI, sarcoidosis) were performed before normalisation so that normalisation statistics were computed exclusively on the retained samples. No outlier removal was applied to validation datasets; PCA plots were generated for visual inspection only.

For GSE56153, the deposited processed series matrix was used as-is under the analysis protocol (median expression = −0.01, classified as processed). Probe-to-symbol collapse on GPL6883 annotation yielded 18,631 unique gene symbols from 24,526 valid-symbol probes; duplicate symbols were resolved using the highest mean-expression probe.

For GSE34608 and GSE81746, the deposited matrices were used as provided. Both are positive and bounded below 20 (GSE34608 median 2.04, range 0.006–13.954; GSE81746 median 0.917, range 0.014– 13.754), consistent with log2/normalised values rather than linear intensities; no additional transformation was applied. Because these cohorts were used only for direction concordance, no AUC was computed for either.

### 2.4 Differential Expression Analysis

#### Microarray datasets (GSE83456 and Bloom)

Differential expression between PTB and HC was assessed using limma [12] with empirical Bayes moderation (eBayes(trend = TRUE, robust = TRUE)) [20]. Covariates were detected automatically from GEO metadata: the GSE83456 model was ∼ 0 + DiseaseStatus + Age + Sex; the Bloom model was ∼ 0 + DiseaseStatus + Sex (Age absent from sub-series metadata). The contrast PTBvsHC = DiseaseStatusPTB − DiseaseStatusHC was applied. *p* values were adjusted using the Benjamini-Hochberg (BH) method to control the false discovery rate (FDR). At FDR < 0.05, GSE83456 yielded 5,513 differentially expressed genes and Bloom yielded 10,174.

#### RNA-seq dataset (GSE107991)

Differential expression was performed using DESeq2 [13] with design ∼ DiseaseStatus (reference level: HC, verified alphabetically via resultsNames()). Log2 fold-change shrinkage was applied using the apeglm method (lfcShrink(type = “apeglm”)) [14]. At FDR < 0.05, 4,758 genes were differentially expressed; fold changes are reported as apeglm-shrunk estimates.

### 2.5 Cross-Cohort Meta-Analysis and Consensus Signature

#### Gene identifier mapping

For GSE107991, Ensembl gene IDs were mapped to HUGO Gene Nomenclature Committee (HGNC) symbols using clusterProfiler::bitr(fromType = “ENSEMBL”, toType = “SYMBOL”, OrgDb = org.Hs.eg.db) [22]. The unmapping rate was 27.4%. Selection was performed on mapped symbols, so all 50 core consensus genes were represented; the unmapped fraction comprised primarily unannotated antisense transcripts, novel transcript loci, and pseudogene classes without stable HGNC symbols. Duplicates were resolved using a keep-first strategy.

#### Common gene universe

A 3-way intersection of mapped gene symbols across GSE83456, Bloom, and GSE107991 yielded a common universe of 15,883 genes, which served as the meta-analysis input and enrichment background.

#### Two-stage gene selection

Gene selection proceeded in two stages. First, a per-study replication criterion was applied: candidate genes were required to achieve BH-adjusted FDR < 0.05 with 100% directional concordance when each dataset was analysed individually (1,959 genes satisfied this criterion). Second, from this per-study-replicated set, meta-analysis effect-size thresholding was applied using a random-effects model fitted by restricted maximum likelihood (REML; metafor::rma(yi, sei, method = “REML”, test = “z”)) [15]: genes were required to achieve pooled |logFC| > 1.5 and random-effects FDR < 0.01, yielding the 50-gene core consensus signature. These 50 genes are the top 2.6% of the 1,959-gene replicated set (97.4th percentile); the same genes lie at the 94.6th percentile of the 932-gene broad consensus and the 99.7th percentile of the 15,883-gene universe. All 50 genes were upregulated in PTB relative to HC.

#### Standard error construction

For the microarray datasets, standard errors fed to metafor::rma() were derived from the limma empirical Bayes posterior variance (s2.post) using the formula sqrt(fit2$s2.post) * fit2$stdev.unscaled[, “PTBvsHC”], checked against the limma author’s guidance on the Bioconductor support channel. For GSE107991, the unshrunken MLE lfcSE from DESeq2::results() was used to ensure meta-analysis weights reflect MLE precision for the RNA-seq contribution.

#### Cross-platform effect-size pooling

Raw log2 fold-changes from limma (microarray, intensity-scale) and apeglm-shrunk log2 fold-changes from DESeq2 (count-scale) were pooled directly rather than converting to Hedges’ g. Both platforms yield log2 fold-changes (same unit), apeglm shrinkage makes the RNA-seq contribution conservative rather than inflated, and the random-effects τ² estimate absorbs residual between-platform variance heterogeneity [16].

#### Heterogeneity and sensitivity analysis

Heterogeneity was assessed using I² per gene; 12 of 50 core genes showed I² ≥ 75% (substantial heterogeneity) but were retained given strong statistical significance, biological plausibility, and the known instability of I² for k = 3 studies [19]. We did not adopt the Hartung–Knapp–Sidik–Jonkman (HKSJ) correction as the primary inference method: with k = 3 studies (2 df), the t-based HKSJ intervals are wide and may be conservative when heterogeneity is large, so they would not provide a usable effect-size threshold for signature construction [17, 18]. The z-test was used for effect-size estimation and thresholding [15]. Signature construction therefore rests on per-study replication, which does not depend on the pooling method: every one of the 50 consensus genes achieves FDR < 0.05 with concordant direction in each of the three individual datasets independently.

### 2.6 Functional Enrichment Analysis

#### Gene set enrichment analysis (GSEA)

GSEA was performed on the full ranked gene lists from all three discovery datasets independently [21]. Ranking metrics differed between platforms: the moderated t-statistic was used for the microarray datasets and the apeglm-shrunk log2 fold-change for GSE107991. Due to this cross-platform incompatibility, Normalised Enrichment Scores (NES) were not pooled across datasets; instead, cross-study concordance was evaluated at the pathway level.

#### Over-representation analysis (ORA)

ORA was performed on the core 50-gene signature (excluding the pseudogene *FCGR1CP*, yielding 49 protein-coding genes as input) against the common universe of 15,883 genes [22]. Gene Ontology Biological Process (GO-BP) results were simplified to reduce redundancy using simplify() with the Wang similarity metric and a cutoff of 0.7.

#### Single-sample gene set enrichment analysis (ssGSEA)

Sample-level enrichment scores were computed using ssGSEA [23] for the 50-gene consensus signature. ssGSEA was applied to the two discovery microarray datasets (GSE83456 and Bloom) and to all validation datasets where AUC was computed. Array coverage differed between platforms, so the number of signature genes represented in each validation cohort is reported with the corresponding estimate: 47 of 50 in GSE28623 and GSE56153.

### 2.7 Protein–Protein Interaction Network Analysis

The 50-gene core consensus signature (excluding the pseudogene *FCGR1CP*; 49 protein-coding genes) was submitted to the STRING protein–protein interaction (PPI) database (v12.0) [24] with a medium confidence threshold (combined score ≥ 400) for *Homo sapiens* (NCBI Taxonomy: 9606). All 49 genes mapped to STRING (100% mapping rate).

Hub genes were identified using the CytoHubba plugin (v0.1) [26] with the Maximal Clique Centrality (MCC) algorithm in Cytoscape (v3.10.4) [25]. Functional module detection was performed using Molecular Complex Detection (MCODE; v2.0.3) [27] with default parameters (degree cutoff = 2, haircut = yes, node score cutoff = 0.2, K-core = 2, max depth = 100).

### 2.8 External Validation

#### Primary validation (GSE28623)

The 50-gene consensus signature was scored in each GSE28623 sample using ssGSEA [23], and the resulting enrichment scores were used to discriminate PTB from HC via receiver operating characteristic (ROC) analysis. Forty-six of the 50 consensus genes matched the GPL4133 annotation by current symbol. The frozen annotation used the obsolete symbol WARS for *WARS1*; after renaming that row, 47 genes were testable. Three genes were absent (*CMPK2*, *FCGR1A*, *FCGR1CP*). The area under the ROC curve (AUC) and 95% confidence intervals were computed using the DeLong method [32]. Statistical significance was assessed using the Wilcoxon rank-sum test.

Individual gene discriminative performance was assessed using per-gene ROC analysis with DeLong confidence intervals. For genes with AUC > 0.90, the Wilcoxon exact test was substituted because the normal approximation degrades near the AUC = 1.0 boundary.

#### Direction concordance (GSE34608 and GSE81746)

For GSE34608 (8 PTB, 18 HC) and GSE81746 (4 PTB, 2 HC), sample sizes were too small for reliable AUC estimation. Direction concordance was computed as the percentage of consensus genes for which the sign of the mean expression difference (PTB − HC) matched the discovery direction.

#### Independent validation (GSE56153)

Forty-five of the 50 consensus genes matched the GPL6883 annotation by current symbol. The frozen annotation used the obsolete symbols WARS and INDO; after renaming them to *WARS1* and *IDO1*, 47 genes were testable (absent: *ETV7*, *CMPK2*, *FCGR1CP*). AUC and DeLong confidence intervals were computed as for the primary validation.

### 2.9 Machine Learning Classifier Development

#### Training configuration

The machine learning classifier was developed using the two microarray discovery datasets (GSE83456 + Bloom post-QC; 219 samples: 70 PTB + 149 HC). The candidate feature space was established upstream from the multi-cohort discovery analysis, comprising the 49 protein-coding consensus genes (*FCGR1CP* pseudogene excluded). The downstream machine learning procedure was designed to perform feature sparsification from this pre-established 49-gene candidate set to a parsimonious classifier. To enable cross-platform application, per-gene z-score normalisation was applied within each dataset independently [16].

#### Feature selection

Least absolute shrinkage and selection operator (LASSO) logistic regression [28] (cv.glmnet(alpha = 1, family = “binomial”)) was used for feature selection with the regularisation parameter set to lambda.1se (the most parsimonious model within one standard error of minimum deviance). Inverse-frequency weights (IFW) were applied during LASSO to prevent majority-class bias; IFW were computed per fold with no pre-computation leakage. The seed was set to 1234.

#### Random forest classifier

A random forest (RF) classifier [30] was trained on the LASSO-selected features with ntree = 500 and mtry = floor(√p). The Synthetic Minority Over-sampling Technique (SMOTE; over_ratio = 1) was applied to the training data only to balance the class distribution [29]. The combination of IFW during feature selection and SMOTE during classification provides a dual imbalance correction, which was empirically validated through sensitivity analysis: IFW+SMOTE, IFW-only, and SMOTE-only variants yielded a maximum AUC difference of 0.001 across 5-fold nested cross-validation.

#### Performance estimation

Model performance was estimated using nested cross-validation: 5-fold stratified outer CV with embedded 10-fold inner CV for LASSO λ tuning. Because the 49 candidate genes were established upstream using the discovery cohorts, this nested cross-validation describes internal performance for the downstream feature sparsification and model training conditional on the pre-established candidate set; it does not represent an unbiased estimate of an end-to-end biomarker-discovery pipeline. SMOTE was applied exclusively to the training portion of each outer fold; validation folds remained untouched.

#### Feature importance

treeshap (v0.4.0) was attempted for SHAP-based feature importance but failed for the 4-feature model (all-NaN output). RF permutation importance (Mean Decrease in Accuracy; importance(type = 1)) was used as the fallback method, which avoids the split-selection bias inherent in Gini impurity measures [31], while feature collinearity was minimised upstream through LASSO feature selection.

### 2.10 PTB versus LTBI Analyses

#### ssGSEA-based discrimination

To evaluate the signature’s ability to discriminate active PTB from LTBI, the 25 LTBI samples excluded from the primary GSE28623 validation were re-incorporated. All 108 samples (46 PTB, 37 HC, 25 LTBI) were jointly re-normalised using normexp background correction (offset = 50), quantile normalisation, GPL4133 probe-to-symbol mapping, removal of invalid symbols, and limma::avereps() probe collapse, yielding 19,749 gene symbols × 108 samples. ssGSEA scores were computed simultaneously for all three groups.

This pipeline differs from the primary validation in two respects: probe collapse uses limma::avereps() rather than max-mean probe selection, and background correction includes an offset = 50 parameter. Both changes are required for joint three-group normalisation. A PTB-versus-HC stability endpoint was computed to confirm that the joint normalisation did not alter the primary validation result (stability criterion: |ΔAUC| ≤ 0.05).

#### Frozen 4-gene RF validation against LTBI

The frozen random forest model (4 genes: *BATF2*, *ANKRD22*, *GBP5* and *EPSTI1*; 500 trees, mtry = 2) was trained on 298 SMOTE-balanced samples (149 PTB, 149 HC) and had an out-of-bag error of 3.02% on that augmented training set. The figure is therefore optimistic and is not an external estimate. The model was applied to GSE28623 without retraining, SMOTE or tuning.

The GSE28623 expression matrix is on a linear intensity scale (median 150.8; range 0.026–633,791), whereas the training matrices are log2-scale and gene-centred. We therefore applied log2 transformation before per-gene z-scoring. This gave a PTB-versus-HC AUC of 0.864 (95% CI 0.785–0.944; n = 83, 46 PTB, 37 HC) and a PTB-versus-LTBI AUC of 0.867 (95% CI 0.784–0.949; n = 71, 46 PTB, 25 LTBI). The fitted model itself was unchanged. The switch from raw-scale to log2-scale z-scoring was made after inspecting the external matrix and recognising the scale mismatch; it was not pre-specified. The superseded raw-scale estimates are recorded in Table S13.

### 2.11 Published Signature Comparators

To contextualise the diagnostic performance of the 50-gene signature, two established blood transcriptomic TB signatures were evaluated on the same primary validation cohort (GSE28623, n = 83: 46 PTB, 37 HC):

**Berry et al. 2010** (393-transcript differentially expressed module) [2]: The official Berry Supplementary Table 3 (393 entries × 6 columns) was used as the source. One entry (Sep-04/SEPT4, Entrez Gene ID 5414) was stored in the supplementary Excel file as an Excel date-like value (38231) and was corrected to SEPT4 (now SEPTIN4) before platform mapping. The mapping chain was: 393 published entries → 312 unique symbols → 278 symbols represented in the GSE28623 matrix. Those 278 symbols account for 338 of the 393 entries (86.0%). The source table contains no per-gene direction column, so the published list cannot by itself specify which genes are up- or down-regulated. We therefore scored the Berry 393-gene set in two ways. First, an unsigned ssGSEA score was computed over all 278 represented genes. Second, a signed score was computed as the difference between the ssGSEA enrichment scores of the up- and down-regulated subsets. Direction was reconstructed from GSE19439, the London training set in which the 393-gene list was derived: 311 of the 312 symbols had a direction estimate there (225 up, 86 down versus HC). Of the 278 symbols represented in GSE28623, 277 had a direction estimate (204 up, 73 down) and entered the signed score. Sign concordance in the GSE19444 test set was 307/311 (98.71%; Spearman ρ of effect sizes 0.891). The signed score is explicitly a reconstruction of direction, not the published Berry score, and is reported alongside the unsigned score rather than replacing it.

**Sweeney et al. 2016** (3-gene diagnostic signature: *GBP5*, *DUSP3*, *KLF2*) [3]: the Sweeney 3-gene score was computed with its published directional TB-score formula (GBP5 + DUSP3)/2 − KLF2, applied to log2-transformed expression values. ssGSEA was not used for this score because it summarises a gene set as a single coordinated enrichment signal and cannot represent genes that move in opposite directions; applying it to a set containing both up- and down-regulated genes would dilute the score. All three Sweeney genes were present on GPL4133.

The Zak et al. 2016 RISK16 signature was not evaluated because it was trained to predict LTBI-to-active-TB progression, not to discriminate active PTB from HC [4].

Pairwise comparisons between signatures were performed using paired DeLong tests on the same 83 samples [32]. The primary Holm family comprised the three comparator contrasts against the 50-gene score (Sweeney 3-gene, unsigned Berry, signed Berry). Mean-z scores for the Berry genes were computed as a sensitivity analysis only and were not entered into the primary correction. Gene-set sizes (50 versus 393 versus 3) and scoring methods (ssGSEA versus directional TB-score) differ, and these results come from a single cohort.

The comparison is also not independent at the level of signature derivation. GSE107991 reanalyses the Berry test set, and two of our discovery datasets overlap Sweeney’s discovery data: Bloom is GSE42834, and GSE107991 reanalyses samples from GSE19444, which belongs to GSE19491 (Table S18b) [2, 3]. GSE28623 was not used to derive either signature, so neither score was fitted to it. It was, however, one of the cohorts in which Sweeney et al. validated their score, so the Sweeney comparison is a re-analysis of a previously reported validation rather than an independent test.

### 2.12 Immune Cell Deconvolution

Immune cell composition was evaluated across the three discovery cohorts using two complementary transcriptomic deconvolution approaches.

**MCPcounter** [34] estimates relative abundance scores for 10 immune and stromal cell populations using transcriptomic marker genes. MCPcounter was selected as the primary method because it does not require a reference signature matrix and is robust across microarray and RNA-seq platforms. MCPcounter gene markers were obtained from the official MCPcounter GitHub repository and cached locally for reproducibility. Input data were log2 expression values for the microarray datasets and log2(TPM + 1) for GSE107991, where TPM denotes transcripts per million.

**CIBERSORTx** [35] was run via the Stanford web portal (https://cibersortx.stanford.edu/) using the LM22 signature matrix (547 genes, 22 immune cell subtypes). CIBERSORTx estimates cell-type proportions by deconvolving bulk expression profiles against a purified cell-type reference. Input data were linear-scale (2^log2) back-transformed expression for the microarray datasets and TPM (linear scale, HUGO symbols) for GSE107991. For GSE107991, TPM values were computed using maximum annotated transcript length per gene (EnsDb.Hsapiens.v86, Ensembl Release 86; 27,882 genes retained with 100% gene-length mapping rate). Ensembl gene IDs were mapped to HGNC symbols via EnsDb.Hsapiens.v86 (27,724 unique symbols retained; 158 duplicate symbols resolved by highest mean expression).

CIBERSORTx portal settings:

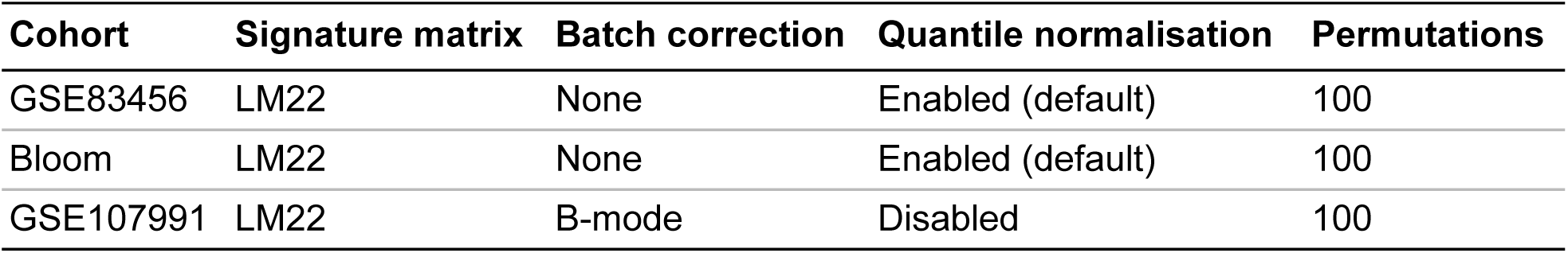

B-mode batch correction was applied to GSE107991 because LM22 was derived from microarray data while GSE107991 is RNA-seq; B-mode corrects cross-platform variation between bulk signature and bulk mixture. S-mode was not used because it requires a single-cell reference matrix. Quantile normalisation was disabled for GSE107991 (RNA-seq TPM input).

Samples with CIBERSORTx deconvolution *p* value > 0.05 were excluded from downstream differential analysis (poor LM22 fit). CIBERSORTx QC pass rates for the microarray cohorts (42.7% for GSE83456; 57.8% for Bloom) reflect platform-specific gene coverage limitations between Illumina HT-12 v4 and the LM22 reference (derived from Affymetrix HGU133A); GSE107991 achieved 100% pass rate with B-mode correction.

### 2.13 Statistical Analysis

#### Differential abundance and correlation testing

For MCPcounter and CIBERSORTx, PTB-versus-HC differences in each cell population were tested using the Wilcoxon rank-sum test. *p* values were adjusted for multiple testing using the Benjamini-Hochberg (BH) method. Cell types with BH-adjusted *p* < 0.05 were reported as significantly differentially abundant. Cross-study concordance was defined as significance (BH *p* < 0.05) in ≥2 of the 3 discovery cohorts.

Spearman rank correlations were computed between each MCPcounter cell-type abundance and the 50-gene ssGSEA enrichment score (per-sample) in the GSE83456 and Bloom cohorts. Correlations with |ρ| > 0.3 and BH-adjusted *p* < 0.05 were reported.

#### Multiple testing correction

All genome-wide analyses (differential expression, meta-analysis, enrichment) applied the BH method for false discovery rate control. The FDR threshold was 0.05 for per-study differential expression and 0.01 for the core consensus signature meta-analysis.

#### ROC analysis

AUC and 95% confidence intervals were computed using the DeLong asymptotic method [32]. Pairwise signature comparisons used paired DeLong tests on identical samples. Statistical significance of ssGSEA score differences between groups was assessed using the Wilcoxon rank-sum test.

#### Software

All analyses were performed in R (v4.6.0; Bioconductor v3.23) [36]. Key packages included limma [12], DESeq2 [13], metafor [15], GSVA [23], clusterProfiler [22], glmnet [28], randomForest [30], pROC [33], and MCPcounter [34]. Network analysis was performed in Cytoscape (v3.10.4) [25] with CytoHubba (v0.1) [26] and MCODE (v2.0.3) [27] plugins. CIBERSORTx [35] was run via the Stanford web portal. STRING v12.0 [24] was accessed via the STRINGdb R package.

## 3. Results

### 3.1 Discovery Cohort Characteristics and Quality Control

Three blood transcriptomic datasets comprising 247 post-quality-control samples (87 PTB + 160 HC) were analysed (Table 1). GSE83456 contributed 103 samples (43 PTB + 60 HC) after removal of 3 outliers (2 PTB + 1 HC) exceeding 3 standard deviations from disease-group centroids. The Bloom dataset (GSE42826 + GSE42830) contributed 116 samples (27 PTB + 89 HC) after removal of 1 HC outlier. GSE107991 contributed 28 samples (17 PTB + 11 HC) after exclusion of 21 LTBI samples and 5 author-flagged outliers. Principal component analysis showed disease-group separation mainly along PC2 in all three discovery cohorts after normalisation, and sub-series analysis of the Bloom dataset confirmed no batch-driven clustering (Figure 1). Quality control plots (PCA, density, boxplot, sample-correlation heatmap) confirmed satisfactory data distributions across all three cohorts (Figures S1–S4).

**Figure 1.**
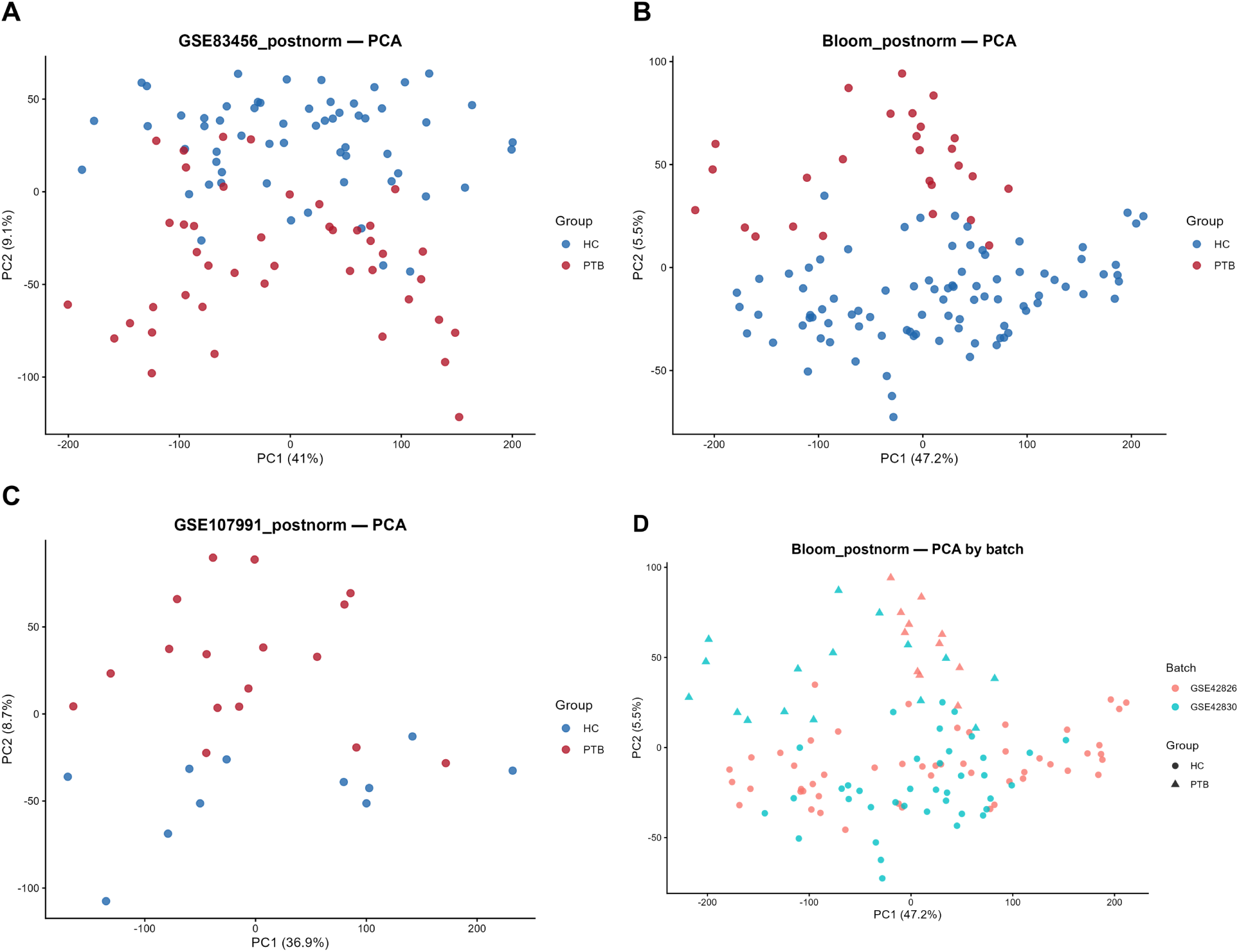
Principal component analysis of the three discovery cohorts. Samples from each discovery cohort were projected onto the first two principal components after normalisation. (A) GSE83456 (n = 103; 43 PTB, 60 HC; Illumina HT-12 v4; PC1 41.0%, PC2 9.1%). (B) Bloom dataset (n = 116; 27 PTB, 89 HC; Illumina HT-12 v4; PC1 47.2%, PC2 5.5%). (C) GSE107991 (n = 28; 17 PTB, 11 HC; RNA-seq; PC1 36.9%, PC2 8.7%). In (A) to (C), samples are coloured by disease status (PTB, red; HC, blue), and disease separation is mainly along PC2 rather than PC1. (D) Bloom dataset coloured by sub-series (GSE42826, pink; GSE42830, teal) with symbol shape denoting disease status (circles, HC; triangles, PTB); the two sub-series do not separate, and PC1 does not track batch origin.

### 3.2 A 50-Gene Consensus Signature Replicates Across Three Discovery Cohorts

A 3-way intersection of mapped gene identifiers gave a common universe of 15,883 symbols. Differential expression at FDR < 0.05 identified 5,513 genes in GSE83456, 10,174 in Bloom and 4,758 in GSE107991. Requiring significance and the same direction in all three studies left 1,959 genes (Table S16).

Meta-analysis then applied FDR < 0.01 and |logFC| > 1.5 and retained 50 genes, the top 2.6% of the 1,959-gene replicated set (97.4th percentile) (Figure 2, Table 2; per-study FDR and direction in Table S1, full meta-analysis in Table S3). All 50 were upregulated in PTB relative to HC, with pooled log2 fold-changes from 1.519 (*ASPHD2*) to 3.867 (*BATF2*). All 50 were a strict subset of the 1,959-gene intersection, so none arose from pooling alone.

**Figure 2.**
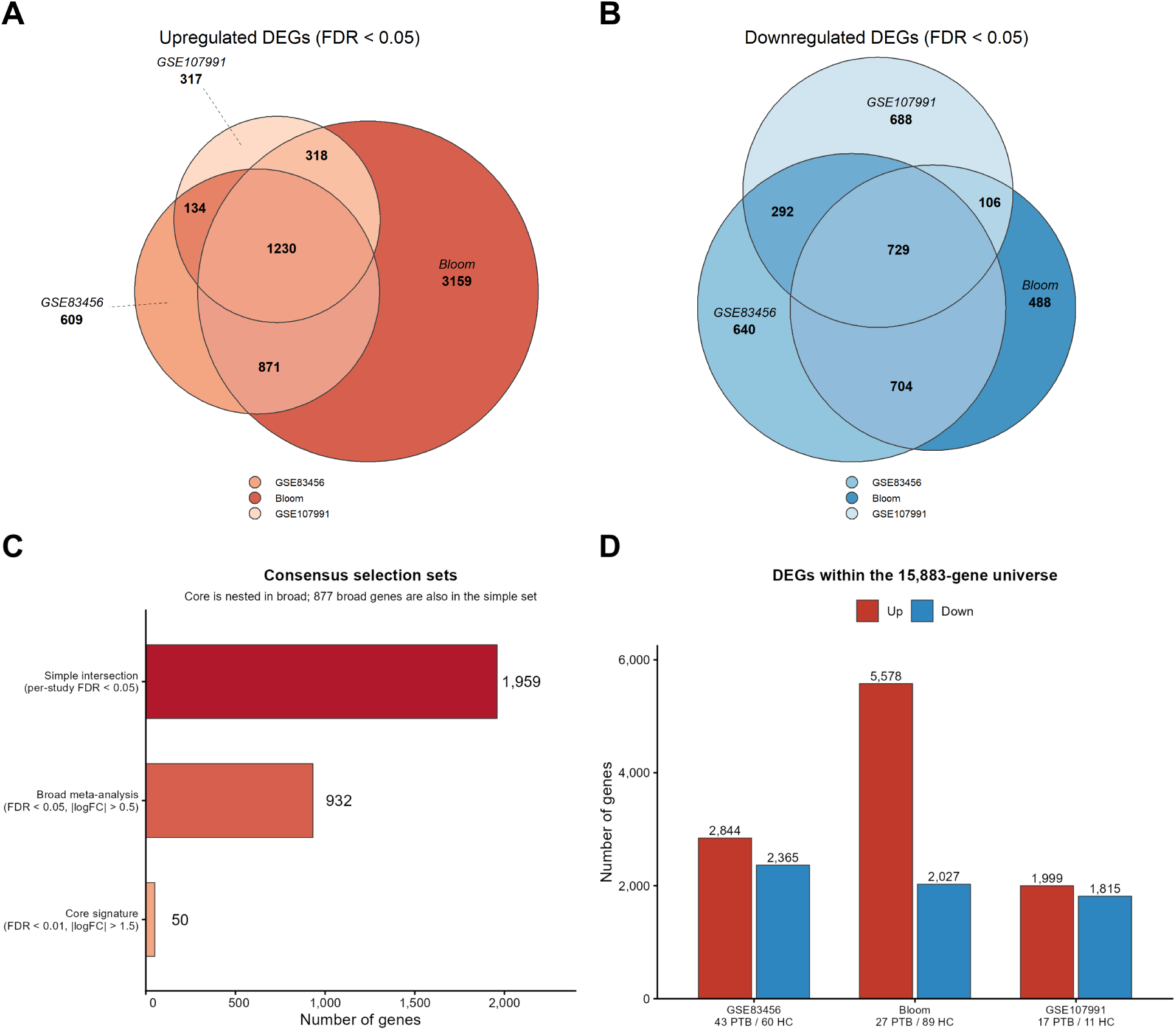
Differential expression overlap and consensus gene selection. Upregulated and downregulated DEG sets were compared across the three discovery cohorts, and consensus tiers were derived from the 15,883-gene universe. (A) Upregulated DEGs (FDR < 0.05); the three-way overlap is 1,230. (B) Downregulated DEGs (FDR < 0.05); the three-way overlap is 729. (C) Consensus tiers: simple intersection (1,959 genes; per-study FDR < 0.05), broad meta-analysis (932 genes; FDR < 0.05, |logFC| > 0.5) and core signature (50 genes; FDR < 0.01, |logFC| > 1.5). The core set is nested inside both the broad tier and the simple intersection, and 877 broad genes are also in the simple set. (D) Numbers of upregulated and downregulated DEGs in each dataset within the 15,883-gene universe. Dataset sample sizes: GSE83456 43 PTB/60 HC; Bloom 27 PTB/89 HC; GSE107991 17 PTB/11 HC.

**TABLE 2.** The 50-Gene Core Consensus Signature for Active Pulmonary Tuberculosis. **Derivation criteria:** Two-stage gene selection combining per-study replication (FDR < 0.05 with 100% directional concordance across GSE83456, Bloom and GSE107991) and random-effects meta-analysis (|logFC| > 1.5, FDR < 0.01 under REML estimation). **Direction:** All 50 genes (100%) were concordantly **upregulated** in PTB versus HC across all three discovery cohorts.

| Gene | Pooled log<sub>2</sub>FC | Meta-analysis FDR | I<sup>2</sup> (%) | Biological module | CytoHubba MCC rank | LASSO selection frequency | GSE83456 FDR | Bloom FDR | GSE107991 FDR |
|---|---|---|---|---|---|---|---|---|---|
| *ANKRD22* | 3.600 | 4.32 × 10<sup>−195</sup> | 0.0% | Other | — | 4/5 (80%) | 4.47 × 10<sup>−27</sup> | 3.29 × 10<sup>−44</sup> | 8.62 × 10<sup>−19</sup> |
| *ASPHD2* | 1.519 | 8.91 × 10<sup>−35</sup> | 55.8% | Other | — | 0/5 (0%) | 7.09 × 10<sup>−24</sup> | 3.62 × 10<sup>−23</sup> | 1.33 × 10<sup>−10</sup> |
| *BATF2* | 3.867 | 4.48 × 10<sup>−223</sup> | 0.0% | Interferon | — | 5/5 (100%) | 3.56 × 10<sup>−25</sup> | 3.43 × 10<sup>−47</sup> | 2.55 × 10<sup>−40</sup> |
| *C1QB* | 3.296 | 3.92 × 10<sup>−11</sup> | **89.9%** | Innate / myeloid | — | 0/5 (0%) | 1.21 × 10<sup>−24</sup> | 1.28 × 10<sup>−23</sup> | 1.67 × 10<sup>−21</sup> |
| *CACNA1E* | 2.113 | 1.84 × 10<sup>−97</sup> | 0.0% | Other | — | 0/5 (0%) | 3.35 × 10<sup>−21</sup> | 1.51 × 10<sup>−26</sup> | 1.05 × 10<sup>−14</sup> |
| *CASP5* | 2.239 | 6.79 × 10<sup>−15</sup> | 74.7% | Innate / myeloid | — | 0/5 (0%) | 3.40 × 10<sup>−17</sup> | 3.21 × 10<sup>−19</sup> | 3.94 × 10<sup>−8</sup> |
| *CEACAM1* | 1.823 | 1.65 × 10<sup>−98</sup> | 0.0% | Innate / myeloid | — | 0/5 (0%) | 1.58 × 10<sup>−23</sup> | 1.36 × 10<sup>−25</sup> | 8.88 × 10<sup>−12</sup> |
| *CLEC4D* | 1.525 | 2.55 × 10<sup>−21</sup> | 42.4% | Innate / myeloid | — | 0/5 (0%) | 7.26 × 10<sup>−14</sup> | 2.56 × 10<sup>−14</sup> | 8.85 × 10<sup>−5</sup> |
| *CMPK2* | 1.595 | 6.03 × 10<sup>−30</sup> | 28.5% | Interferon | — | 0/5 (0%) | 1.50 × 10<sup>−14</sup> | 9.68 × 10<sup>−15</sup> | 2.11 × 10<sup>−6</sup> |
| *CXCL10* | 2.115 | 2.15 × 10<sup>−30</sup> | 43.3% | Interferon | — | 0/5 (0%) | 6.54 × 10<sup>−14</sup> | 1.07 × 10<sup>−16</sup> | 1.59 × 10<sup>−4</sup> |
| *EPSTI1* | 2.734 | 7.80 × 10<sup>−32</sup> | **77.5%** | Interferon | 7 (tie) | 3/5 (60%) | 7.22 × 10<sup>−27</sup> | 2.82 × 10<sup>−35</sup> | 4.73 × 10<sup>−20</sup> |
| *ETV7* | 2.397 | 1.23 × 10<sup>−5</sup> | **96.3%** | Interferon | — | 1/5 (20%) | 2.98 × 10<sup>−29</sup> | 1.99 × 10<sup>−26</sup> | 9.25 × 10<sup>−18</sup> |
| FBXO6 | 2.043 | 2.31 × 10<sup>−141</sup> | 0.0% | Innate / myeloid | — | 0/5 (0%) | 1.64 × 10<sup>−26</sup> | 9.63 × 10<sup>−33</sup> | 1.08 × 10<sup>−20</sup> |
| *FCGR1A* | 3.499 | 1.87 × 10<sup>−172</sup> | 0.0% | Interferon | — | 2/5 (40%) | 6.02 × 10<sup>−25</sup> | 1.13 × 10<sup>−40</sup> | 8.60 × 10<sup>−22</sup> |
| *FCGR1CP* | 3.623 | 8.69 × 10<sup>−100</sup> | 29.1% | Innate / myeloid | — | n/a | 9.08 × 10<sup>−23</sup> | 4.07 × 10<sup>−34</sup> | 1.91 × 10<sup>−33</sup> |
| *GBP1* | 2.642 | 1.00 × 10<sup>−52</sup> | 60.6% | Interferon | 1 (tie) | 0/5 (0%) | 3.89 × 10<sup>−21</sup> | 2.18 × 10<sup>−39</sup> | 1.31 × 10<sup>−25</sup> |
| *GBP4* | 1.848 | 6.10 × 10<sup>−116</sup> | 0.0% | Interferon | — | 1/5 (20%) | 1.93 × 10<sup>−23</sup> | 2.48 × 10<sup>−30</sup> | 6.53 × 10<sup>−16</sup> |
| *GBP5* | 2.907 | 3.60 × 10<sup>−49</sup> | 71.9% | Interferon | — | 5/5 (100%) | 1.48 × 10<sup>−24</sup> | 3.43 × 10<sup>−47</sup> | 6.49 × 10<sup>−33</sup> |
| *GBP6* | 3.065 | 8.50 × 10<sup>−7</sup> | **96.1%** | Interferon | — | 0/5 (0%) | 6.25 × 10<sup>−22</sup> | 9.25 × 10<sup>−44</sup> | 8.65 × 10<sup>−23</sup> |
| HERC5 | 1.775 | 1.93 × 10<sup>−59</sup> | 0.0% | Interferon | — | 0/5 (0%) | 3.37 × 10<sup>−18</sup> | 4.64 × 10<sup>−19</sup> | 1.17 × 10<sup>−3</sup> |
| *IDO1* | 1.539 | 1.33 × 10<sup>−7</sup> | **79.5%** | Interferon | — | 0/5 (0%) | 1.57 × 10<sup>−7</sup> | 2.64 × 10<sup>−8</sup> | 2.36 × 10<sup>−2</sup> |
| *IFI27* | 2.605 | 1.37 × 10<sup>−9</sup> | 67.7% | Interferon | — | 0/5 (0%) | 1.73 × 10<sup>−8</sup> | 9.46 × 10<sup>−7</sup> | 1.58 × 10<sup>−6</sup> |
| *IFI35* | 1.650 | 8.01 × 10<sup>−67</sup> | 39.7% | Interferon | 1 (tie) | 0/5 (0%) | 1.06 × 10<sup>−24</sup> | 2.17 × 10<sup>−33</sup> | 1.43 × 10<sup>−12</sup> |
| *IFI44* | 2.269 | 4.30 × 10<sup>−66</sup> | 19.7% | Interferon | — | 0/5 (0%) | 1.16 × 10<sup>−22</sup> | 8.91 × 10<sup>−25</sup> | 1.04 × 10<sup>−6</sup> |
| *IFI44L* | 2.376 | 1.63 × 10<sup>−68</sup> | 0.0% | Interferon | — | 0/5 (0%) | 2.59 × 10<sup>−20</sup> | 8.34 × 10<sup>−19</sup> | 1.66 × 10<sup>−5</sup> |
| *IFIT1* | 1.906 | 2.17 × 10<sup>−41</sup> | 0.0% | Interferon | — | 0/5 (0%) | 8.60 × 10<sup>−15</sup> | 4.57 × 10<sup>−14</sup> | 5.43 × 10<sup>−3</sup> |
| *IFIT3* | 2.516 | 9.61 × 10<sup>−18</sup> | 74.5% | Interferon | 4 (tie) | 0/5 (0%) | 9.08 × 10<sup>−23</sup> | 3.60 × 10<sup>−21</sup> | 3.32 × 10<sup>−7</sup> |
| *ISG15* | 2.318 | 3.05 × 10<sup>−57</sup> | 0.0% | Interferon | 4 (tie) | 0/5 (0%) | 9.14 × 10<sup>−17</sup> | 3.64 × 10<sup>−17</sup> | 5.78 × 10<sup>−7</sup> |
| LAP3 | 1.801 | 6.20 × 10<sup>−12</sup> | **85.3%** | Interferon | — | 0/5 (0%) | 2.08 × 10<sup>−16</sup> | 4.85 × 10<sup>−29</sup> | 5.18 × 10<sup>−12</sup> |
| *LHFPL2* | 1.842 | 1.56 × 10<sup>−113</sup> | 0.0% | Other | — | 0/5 (0%) | 7.82 × 10<sup>−26</sup> | 8.72 × 10<sup>−27</sup> | 1.00 × 10<sup>−15</sup> |
| *MYL9* | 1.610 | 2.60 × 10<sup>−13</sup> | 56.5% | Other | — | 0/5 (0%) | 2.30 × 10<sup>−11</sup> | 6.63 × 10<sup>−13</sup> | 2.62 × 10<sup>−4</sup> |
| *OAS3* | 1.662 | 4.10 × 10<sup>−4</sup> | **90.7%** | Interferon | 7 (tie) | 0/5 (0%) | 2.36 × 10<sup>−8</sup> | 2.61 × 10<sup>−19</sup> | 1.36 × 10<sup>−5</sup> |
| OASL | 1.618 | 1.81 × 10<sup>−12</sup> | **75.2%** | Interferon | — | 0/5 (0%) | 6.43 × 10<sup>−21</sup> | 6.98 × 10<sup>−12</sup> | 1.48 × 10<sup>−7</sup> |
| *P2RY14* | 1.700 | 1.50 × 10<sup>−57</sup> | 15.2% | Interferon | — | 0/5 (0%) | 1.77 × 10<sup>−13</sup> | 1.93 × 10<sup>−28</sup> | 7.93 × 10<sup>−8</sup> |
| PARP14 | 1.612 | 1.91 × 10<sup>−127</sup> | 0.0% | Interferon | — | 0/5 (0%) | 7.22 × 10<sup>−27</sup> | 1.97 × 10<sup>−31</sup> | 4.86 × 10<sup>−11</sup> |
| PLSCR1 | 1.752 | 3.26 × 10<sup>−52</sup> | 0.0% | Interferon | — | 0/5 (0%) | 1.28 × 10<sup>−15</sup> | 4.43 × 10<sup>−16</sup> | 1.08 × 10<sup>−7</sup> |
| *PSTPIP2* | 1.681 | 9.32 × 10<sup>−31</sup> | 67.7% | Innate / myeloid | — | 0/5 (0%) | 1.89 × 10<sup>−22</sup> | 8.03 × 10<sup>−30</sup> | 6.71 × 10<sup>−14</sup> |
| *RSAD2* | 2.989 | 4.57 × 10<sup>−81</sup> | 0.0% | Interferon | 4 (tie) | 0/5 (0%) | 1.10 × 10<sup>−21</sup> | 3.05 × 10<sup>−22</sup> | 1.37 × 10<sup>−6</sup> |
| RTP4 | 2.369 | 5.47 × 10<sup>−122</sup> | 13.4% | Interferon | — | 0/5 (0%) | 1.21 × 10<sup>−24</sup> | 5.66 × 10<sup>−35</sup> | 5.84 × 10<sup>−24</sup> |
| *SAMD9L* | 1.773 | 1.69 × 10<sup>−49</sup> | 55.3% | Interferon | 7 (tie) | 0/5 (0%) | 8.27 × 10<sup>−26</sup> | 3.67 × 10<sup>−33</sup> | 3.34 × 10<sup>−12</sup> |
| SLAMF8 | 1.527 | 2.12 × 10<sup>−6</sup> | **90.3%** | Innate / myeloid | — | 0/5 (0%) | 1.43 × 10<sup>−21</sup> | 1.14 × 10<sup>−11</sup> | 4.78 × 10<sup>−12</sup> |
| *SLC6A12* | 1.690 | 5.30 × 10<sup>−74</sup> | 0.0% | Other | — | 0/5 (0%) | 1.36 × 10<sup>−17</sup> | 1.55 × 10<sup>−21</sup> | 3.87 × 10<sup>−13</sup> |
| *SOCS1* | 1.628 | 2.30 × 10<sup>−78</sup> | 0.0% | Interferon | — | 0/5 (0%) | 3.10 × 10<sup>−23</sup> | 5.15 × 10<sup>−19</sup> | 2.08 × 10<sup>−8</sup> |
| *STAT1* | 1.553 | 4.65 × 10<sup>−17</sup> | **86.7%** | Interferon | 1 (tie) | 0/5 (0%) | 1.13 × 10<sup>−23</sup> | 3.02 × 10<sup>−32</sup> | 5.84 × 10<sup>−24</sup> |
| *TCN2* | 1.534 | 3.27 × 10<sup>−7</sup> | **88.1%** | Other | — | 0/5 (0%) | 7.20 × 10<sup>−14</sup> | 2.25 × 10<sup>−13</sup> | 1.01 × 10<sup>−17</sup> |
| *TIMM10* | 2.135 | 3.62 × 10<sup>−72</sup> | 0.0% | Other | — | 0/5 (0%) | 5.49 × 10<sup>−20</sup> | 5.84 × 10<sup>−17</sup> | 6.08 × 10<sup>−14</sup> |
| TRIM22 | 1.595 | 1.15 × 10<sup>−117</sup> | 0.0% | Interferon | — | 0/5 (0%) | 1.70 × 10<sup>−22</sup> | 1.00 × 10<sup>−31</sup> | 9.88 × 10<sup>−16</sup> |
| VAMP5 | 1.812 | 4.37 × 10<sup>−92</sup> | 18.4% | Interferon | — | 0/5 (0%) | 2.58 × 10<sup>−25</sup> | 1.89 × 10<sup>−30</sup> | 2.45 × 10<sup>−14</sup> |
| *WARS1* | 1.816 | 7.39 × 10<sup>−15</sup> | **82.4%** | Interferon | — | 0/5 (0%) | 2.48 × 10<sup>−19</sup> | 2.83 × 10<sup>−32</sup> | 9.77 × 10<sup>−9</sup> |
| *XAF1* | 1.633 | 1.92 × 10<sup>−33</sup> | 53.1% | Interferon | 7 (tie) | 0/5 (0%) | 3.95 × 10<sup>−16</sup> | 1.17 × 10<sup>−30</sup> | 5.17 × 10<sup>−8</sup> |

**Notes**
1. **Gene symbol:** Official HGNC approved symbol. *FCGR1CP* is a transcribed pseudogene; the remaining 49 genes are protein-coding.
2. **Pooled log<sub>2</sub>FC:** Meta-analysis pooled log<sub>2</sub> fold-change under a random-effects model with restricted maximum likelihood (REML) estimation.
3. **Meta-analysis FDR:** Benjamini-Hochberg false discovery rate across all 15,883 genes in the common discovery universe.
4. **I<sup>2</sup> (%):** Percentage of total variation across studies due to heterogeneity rather than chance. Values at or above 75% are shown in bold; twelve genes reach this level and were retained because I<sup>2</sup> is unstable when k = 3.
5. **Biological module:** Interferon response denotes membership of the MSigDB Hallmark interferon-α or interferon-γ set (29 of the 34 genes) plus five established interferon-stimulated genes not captured by those sets (*ETV7*, *GBP1*, *GBP5*, HERC5 and TRIM22). Innate / myeloid denotes innate immune and myeloid activation (8 genes); Other covers the remainder (8 genes). The gene-level annotation and the Hallmark membership check are in Table S19.
6. **CytoHubba MCC rank:** Topological centrality rank in the STRING PPI network (combined score ≥ 0.400) using the Maximal Clique Centrality algorithm in Cytoscape.
7. **LASSO selection frequency:** Number of outer folds (of 5) in nested cross-validation in which the gene was retained at λ.1se. *BATF2*, *GBP5*, *ANKRD22* and *EPSTI1* form the frozen 4-gene classifier.
8. **Per-study FDR:** Benjamini-Hochberg adjusted *p* values from individual cohort differential expression analyses (limma for GSE83456 and Bloom; DESeq2 apeglm for GSE107991). The maximum individual FDR across all 50 genes is 2.36 × 10<sup>−2</sup> (GSE107991, *IDO1*).

Every one of the 50 genes reached FDR < 0.05 in each dataset separately; the weakest was 2.36 × 10⁻² in GSE107991. Direction concordance was 100%: all 50 were upregulated in all three studies. The 50-gene set is therefore replicated study by study, with effect sizes estimated by meta-analysis.

Twelve of the 50 genes (24%) had I² ≥ 75%, among them *STAT1* (86.7%), *ETV7* (96.3%) and *GBP6* (96.1%). We kept them because they were strongly significant and biologically plausible, and because I² is unstable when k = 3 [19]. Figure S6 shows within-discovery ssGSEA scores for GSE83456 and Bloom. These are apparent (resubstitution) values and are not used as validation estimates.

### 3.3 Functional Enrichment Shows Interferon Activation with Ribosomal and Translational Suppression

GSEA run separately in each discovery dataset identified 969 GO-BP terms at FDR < 0.05 in GSE83456, 1,419 in Bloom and 29 in GSE107991 (Figure 3, Table S12). Two programmes appeared in all three. The activated one was dominated by interferon signalling, innate immune activation and viral-response terms; the suppressed one by cytoplasmic translation, ribosome biogenesis and rRNA processing. Of the 858 GO-BP terms significant in at least two datasets, all 858 were directionally concordant, and 24 were significant in all three. Among the wider set of 1,392 terms tested in at least two datasets, 1,383 (99.35%) had concordant normalised enrichment scores. The 9 exceptions were not significant in two or more datasets and touched neither interferon nor viral-response pathways; they included mitochondrial electron transport, T-cell receptor signalling and microvillus assembly.

**Figure 3.**
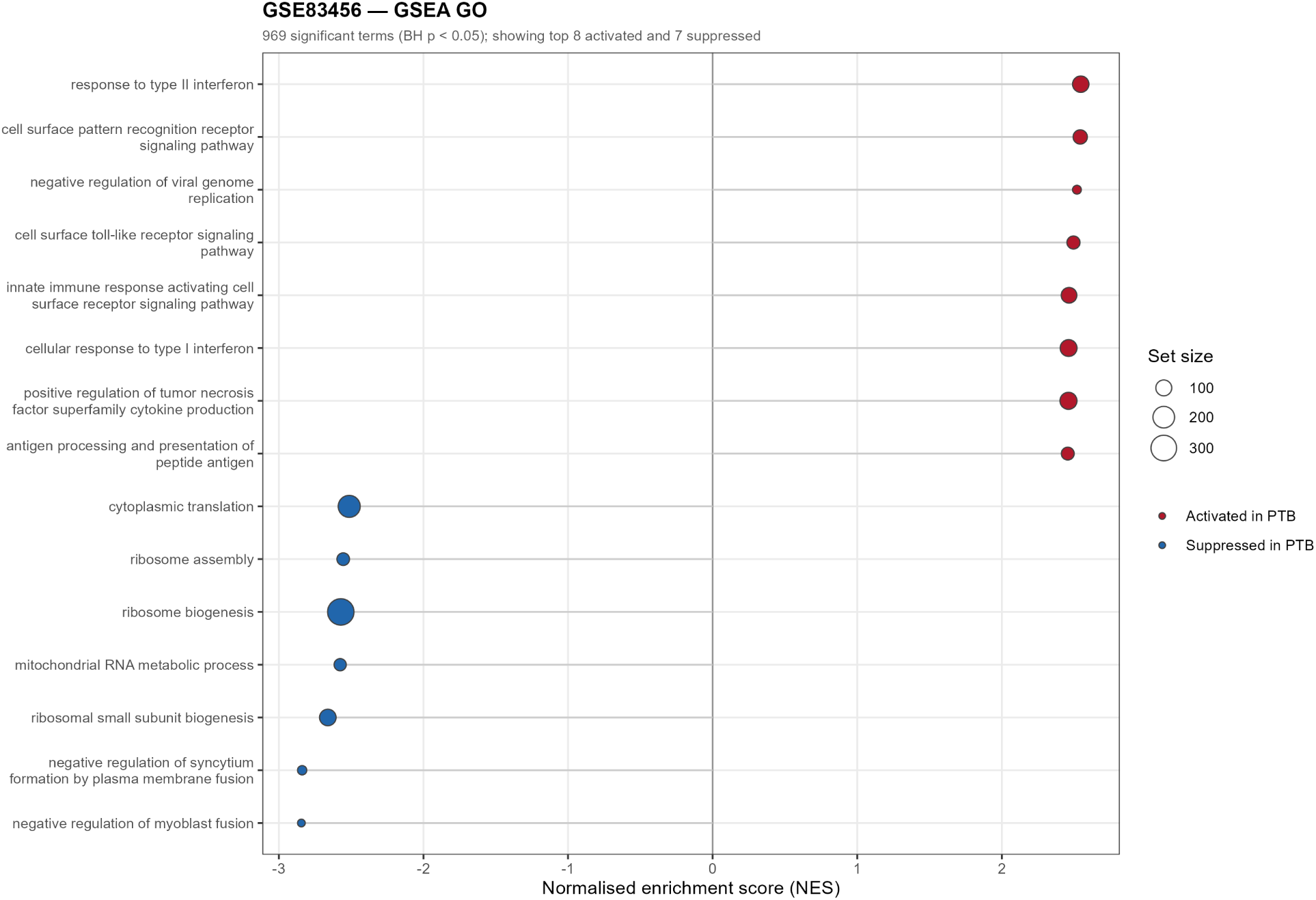
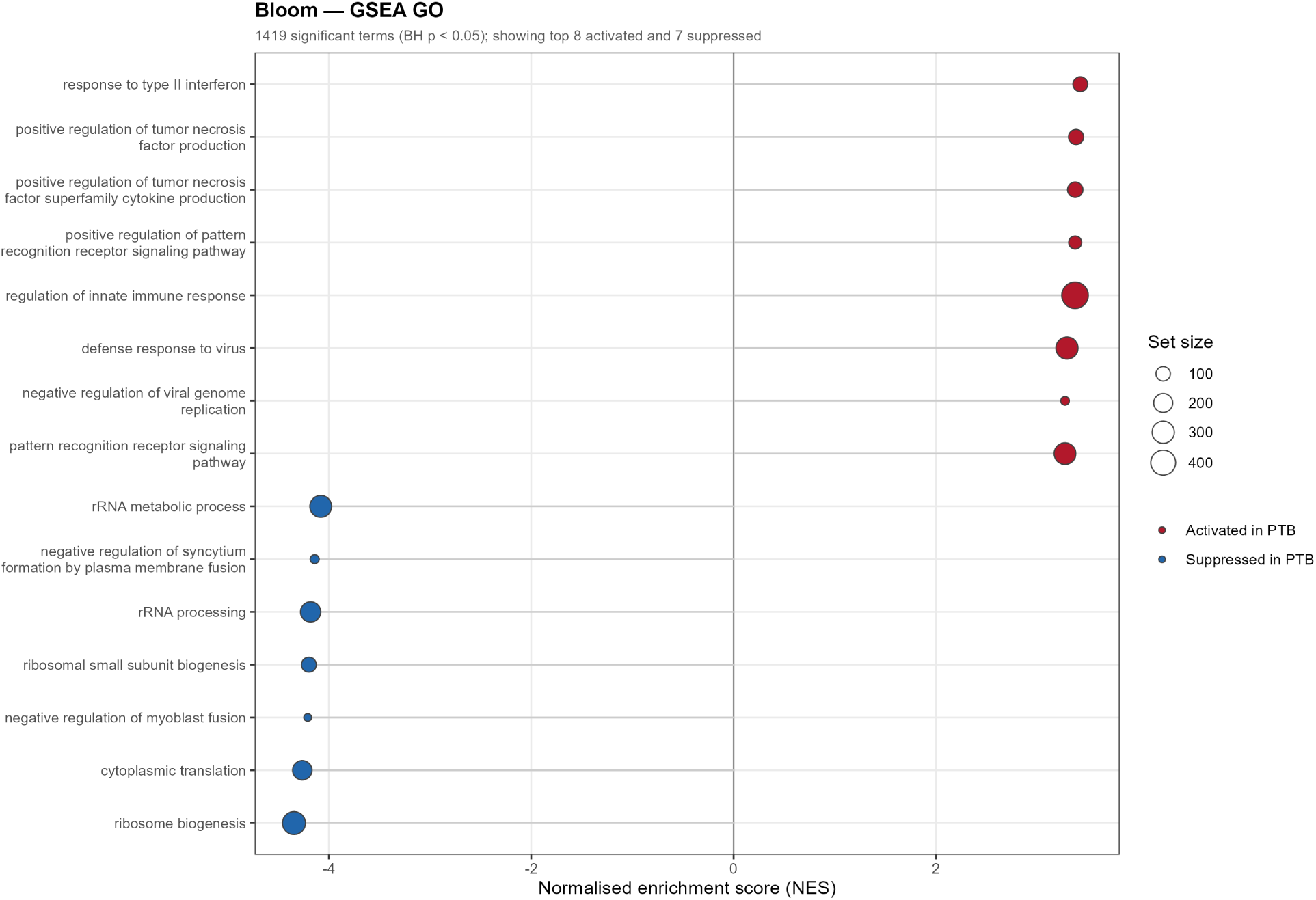
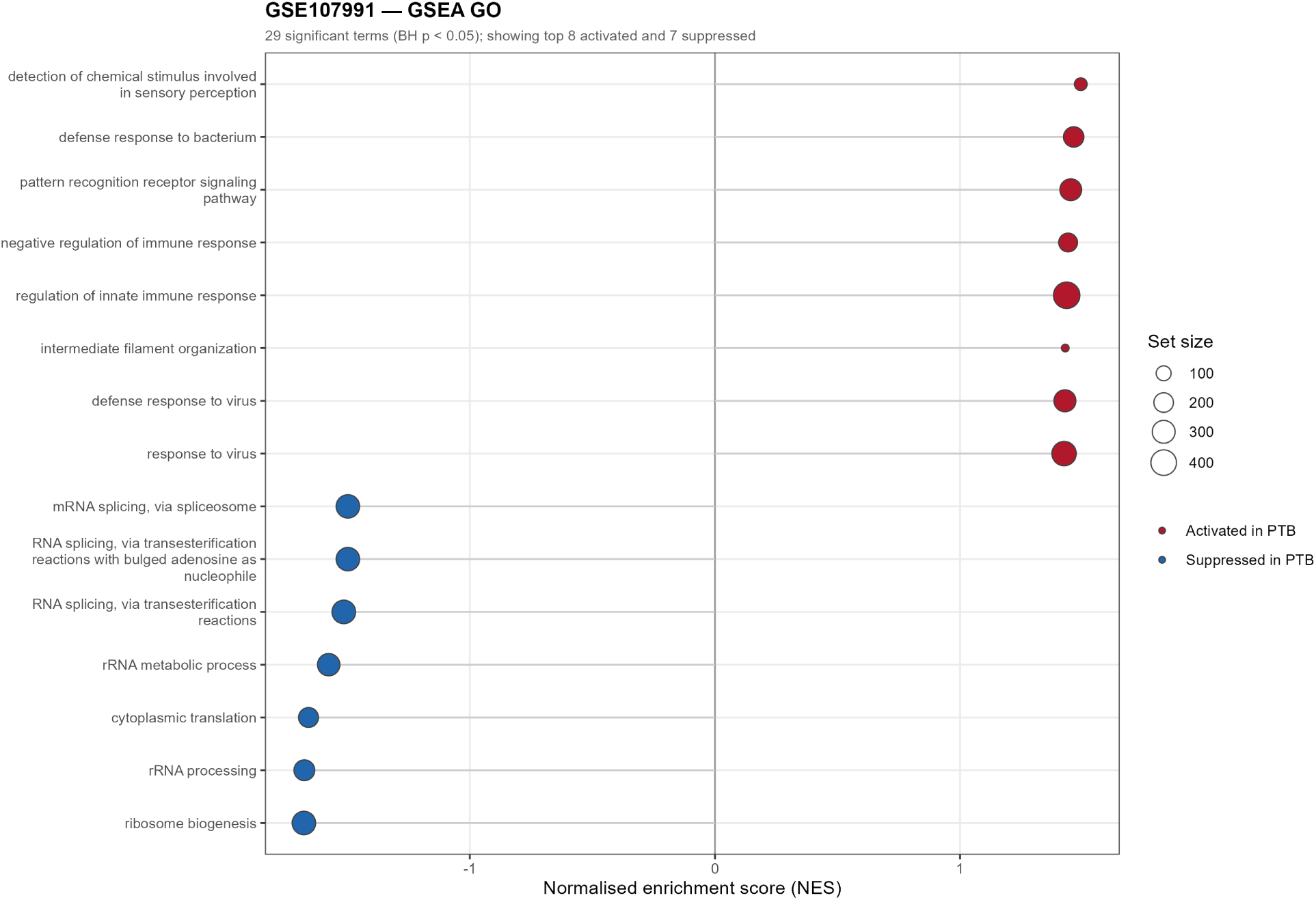
Functional enrichment. Gene set enrichment analysis (GSEA) of GO Biological Process terms in each discovery cohort. (A) GSE83456 (969 significant terms). (B) Bloom (1,419). (C) GSE107991 (29; one additional significant term had undefined NES and is not displayed). Each panel shows the eight most strongly activated and seven most strongly suppressed terms among those at BH FDR < 0.05; the x-axis is normalised enrichment score, dot size is gene-set size, and colour denotes direction in PTB. Interferon signalling, innate immune activation, and viral-defence terms are activated in all three datasets, while cytoplasmic translation, ribosome biogenesis, and rRNA processing are suppressed. (B) Bloom dataset: 1,419 significant GO-BP terms at BH FDR < 0.05. The eight most strongly activated and seven most strongly suppressed terms are shown; dot size is gene-set size and colour denotes direction in PTB. Same conventions as panel (A). (C) GSE107991: 29 significant GO-BP terms at BH FDR < 0.05. The eight most strongly activated and seven most strongly suppressed terms are shown; dot size is gene-set size and colour denotes direction in PTB. Same conventions as panel (A).

ORA used the 49 protein-coding consensus genes, with the pseudogene *FCGR1CP* excluded, against the 15,883-gene background and returned 33 GO-BP terms, mostly immune or antiviral, and 2 KEGG pathways (Figure 4; unsimplified GO output in Figure S9). Figure S5 shows KEGG-pathway GSEA for each cohort. Ribosome and ribosome-biogenesis pathways were suppressed; phagocytosis, lysosome, NOD-like receptor and TNF signalling were activated. Only *FCGR1A* from the signature maps to the “Tuberculosis” KEGG pathway (hsa05152), so that pathway was not significant.

**Figure 4.**
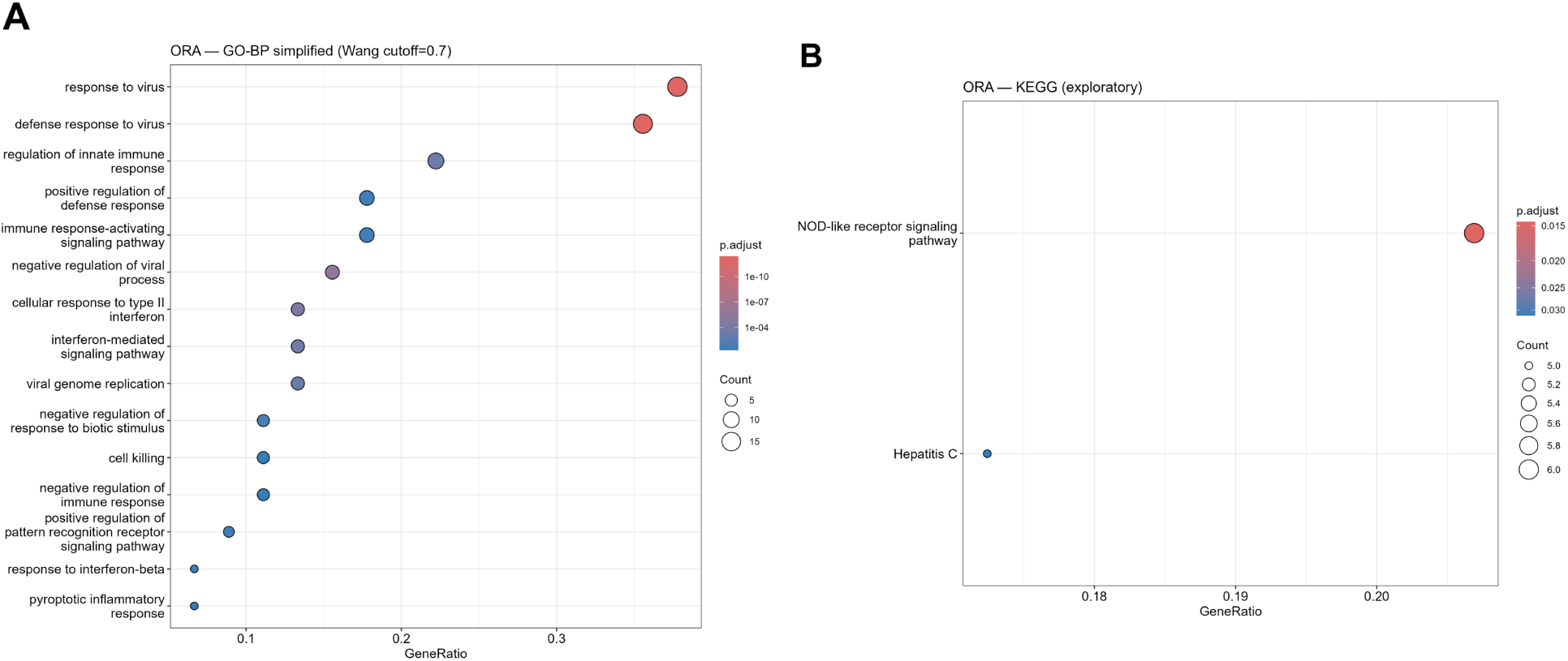
Over-representation analysis of the 50-gene signature. The 49 protein-coding consensus genes (*FCGR1CP* pseudogene excluded) were tested against the 15,883-gene background. (A) Simplified GO Biological Process enrichment; the 15 most significant terms are displayed, drawn from 33 significant terms in the full output. (B) KEGG pathway enrichment (2 significant pathways). Only *FCGR1A* maps to the “Tuberculosis” KEGG pathway (hsa05152), which was therefore not significant.

Annotated against the MSigDB Hallmark interferon-α and interferon-γ sets, with five additional established interferon-stimulated genes added under the fixed rule described in Table 2 note 5, the 50 genes fall into three modules: interferon response (34 genes, 29 of which are in the Hallmark union, plus *ETV7*, *GBP1*, *GBP5*, HERC5 and TRIM22), innate immune/myeloid activation (8, including *C1QB*, *CEACAM1*, *CLEC4D* and *PSTPIP2*) and other (8, including *ANKRD22*, *CACNA1E* and *LHFPL2*; Table S19). That every gene was upregulated fits the dominant interferon module of Berry et al. (2010) [2], whose 393-gene list was not filtered on direction; in GSE19439, 86 of the 311 measurable Berry genes are down-regulated in PTB. Down-regulated pathways such as lymphocyte signalling have smaller effect sizes. They appear in the broad consensus (932 genes at |logFC| > 0.5) but not in the core signature (Table S2).

### 3.4 A Dense Interferon Signalling Module in the Protein–Protein Interaction Network

STRING analysis of the 49 protein-coding consensus genes at medium confidence (≥ 0.4) found 363 edges among 41 connected nodes and 8 isolated nodes (*CACNA1E*, *TIMM10*, *TCN2*, *ASPHD2*, *SLC6A12*, *P2RY14*, *MYL9*, *LHFPL2*) (Figure 5). Seven of the 8 isolates belong to the other module and have no established interactions with the core interferon machinery.

**Figure 5.**
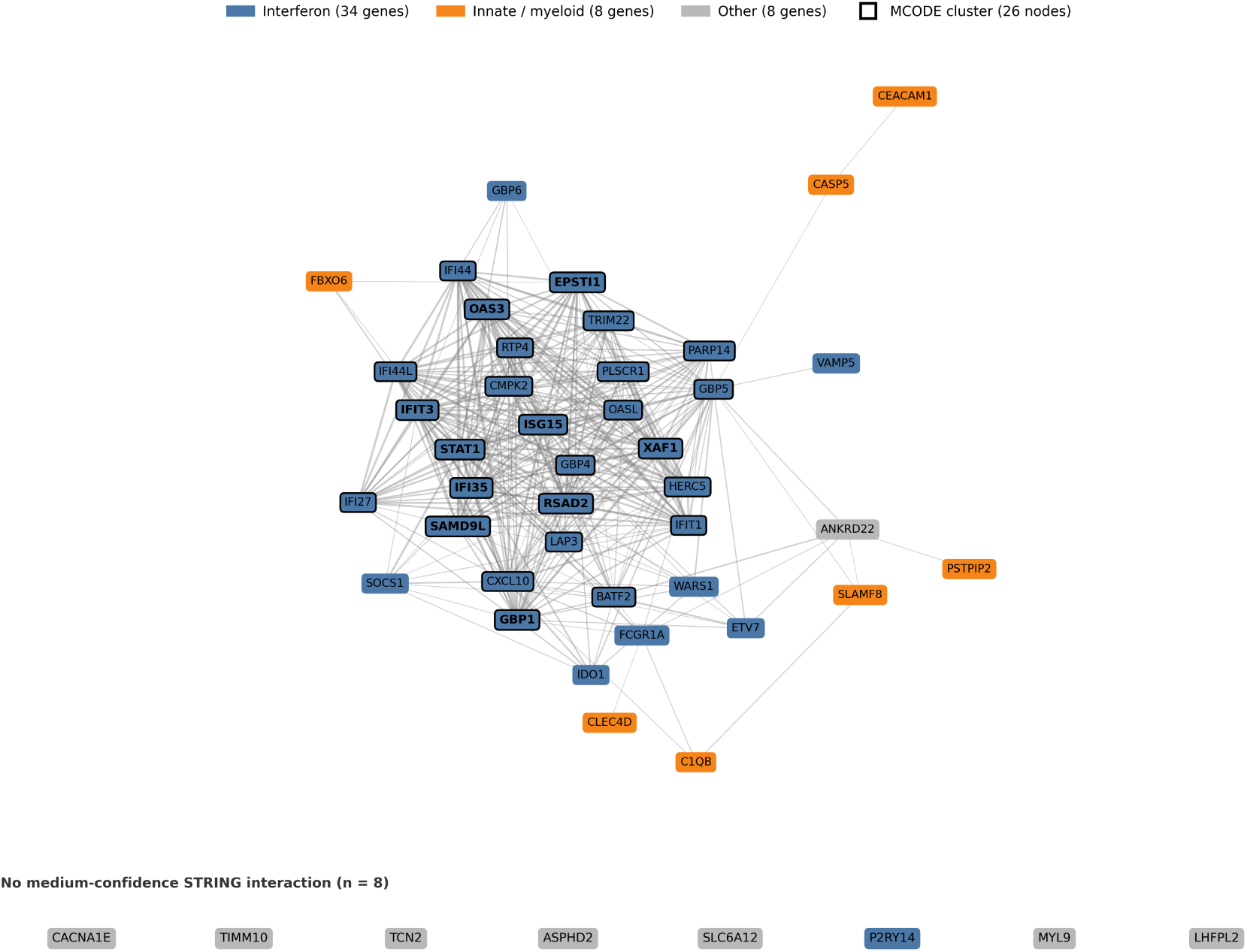
Protein–protein interaction network. STRING v12.0 (medium confidence ≥ 0.4) yielded 363 interactions among 41 of the 49 protein-coding consensus genes; the 8 genes with no medium-confidence interaction are shown separately at the bottom. Nodes are coloured by module (interferon, blue; innate/myeloid, orange; other, grey), and the MCODE cluster (26 nodes, score 24.480) is outlined in black. The CytoHubba MCC top 10 are shown in bold: *STAT1*, *IFI35*, *GBP1*, *IFIT3*, *RSAD2*, *ISG15*, *XAF1*, *SAMD9L*, *EPSTI1* and *OAS3*.

CytoHubba MCC returned 10 hub genes, all interferon-stimulated: *STAT1*, *IFI35* and *GBP1* tied at the top, followed by *IFIT3*, *RSAD2*, *ISG15*, *XAF1*, *SAMD9L*, *EPSTI1* and *OAS3*. *STAT1*, the master transcription factor of interferon signalling, headed the list.

MCODE found one dense module of 26 nodes (63% of the 41 connected genes), with 306 edges and a score of 24.480. It contained the interferon-driven core: *STAT1*, *ISG15*, *RSAD2*, *GBP1*, *GBP5*, *IFIT1*, *IFIT3* and *CXCL10*. The remaining 15 connected nodes lay outside the dense module.

### 3.5 Cross-Platform External Validation

#### Primary validation (GSE28623, The Gambia)

We evaluated the 50-gene signature in the external Gambian cohort (GSE28623; 46 PTB + 37 HC; Agilent GPL4133), which was held out entirely: no gene selection, threshold or model choice used validation data. Forty-seven of the 50 genes were represented on the platform after alias correction. ssGSEA scores discriminated PTB from HC with an AUC of 0.830 (95% CI 0.743–0.917; Wilcoxon *p* = 5.912 × 10⁻⁸) (Figure 6, Table 3).

**Figure 6.**
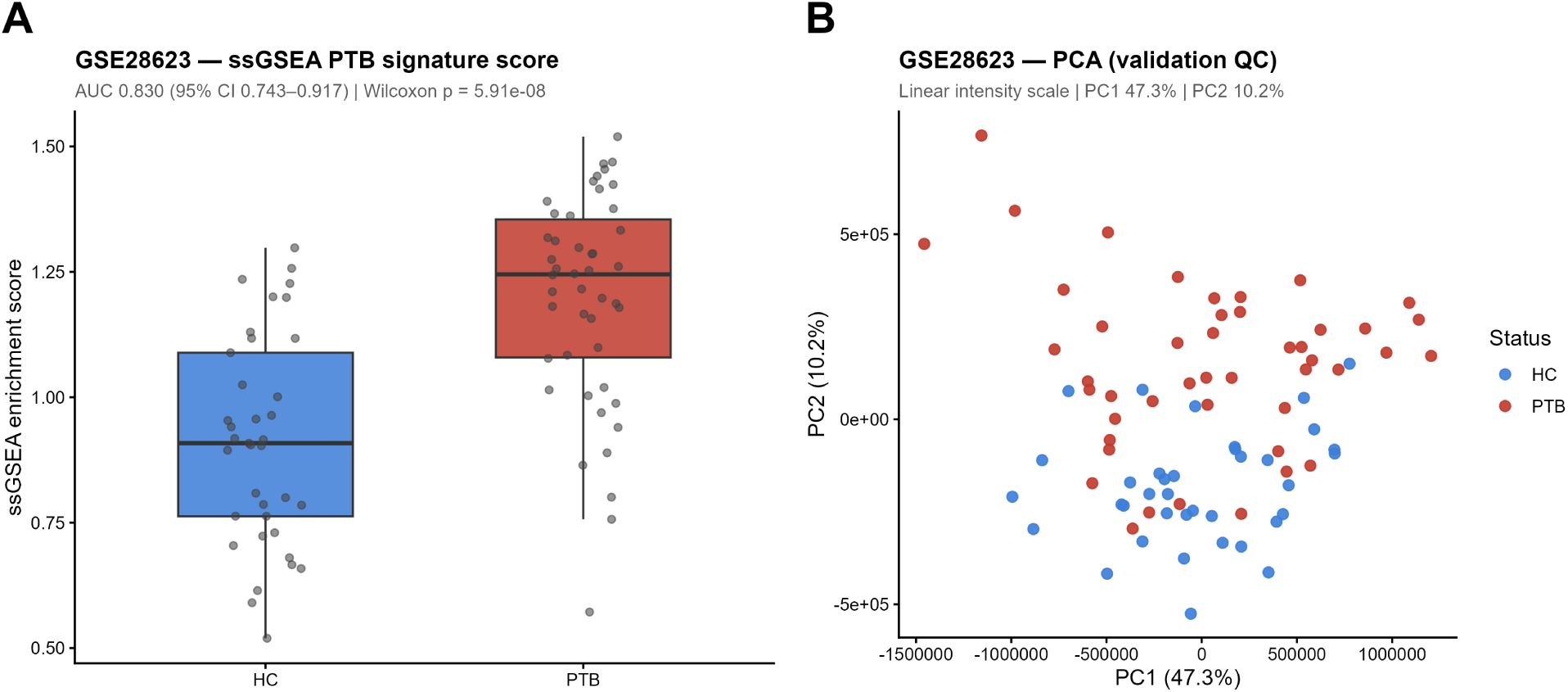
External validation. The 50-gene consensus signature was tested in the Gambian cohort (GSE28623; 46 PTB, 37 HC; Agilent GPL4133). (A) Boxplot of ssGSEA enrichment scores by disease status. The signature discriminated PTB from HC with AUC = 0.830 (95% CI 0.743–0.917; exact Wilcoxon p = 5.912 × 10⁻⁸). (B) PCA of the GSE28623 validation cohort on the linear intensity scale, coloured by disease status; PC1 explains 47.3% and PC2 10.2%, and disease separation is mainly along PC2.

**TABLE 3.** Consolidated Validation Performance Summary.

| **Analysis** | **Cohort** | **Score / model** | **AUC** | **95% CI** | **Wilcoxon *p*** | **Key note** |
|---|---|---|---|---|---|---|
| **Primary 50-gene score** | GSE28623 (46 PTB + 37 HC) | 50-gene ssGSEA (47 represented) | **0.830** | 0.743–0.917 | 5.912 × 10<sup>−8</sup> | Primary external validation anchor |
| **Joint normalisation, PTB versus HC** | GSE28623 (108-sample joint normalisation) | 50-gene ssGSEA | **0.831** | 0.745–0.918 | 5.223 × 10<sup>−8</sup> | ΔAUC +0.0012; preprocessing stability |
| **Joint normalisation, PTB versus LTBI** | GSE28623 (46 PTB + 25 LTBI) | 50-gene ssGSEA | **0.848** | 0.756–0.940 | 3.238 × 10<sup>−7</sup> | Active versus latent TB discrimination |
| **4-gene classifier, external** | GSE28623 (46 PTB + 37 HC) | 4-gene RF (frozen; log2 + z) | **0.864** | 0.785–0.944 | 9.77 × 10<sup>−10</sup> | Primary external estimate |
| **4-gene classifier, LTBI** | GSE28623 (46 PTB + 25 LTBI) | 4-gene RF (frozen; log2 + z) | **0.867** | 0.784–0.949 | 4.164 × 10<sup>−8</sup> | Active versus latent TB discrimination |
| **Sweeney 3-gene score** | GSE28623 (46 PTB + 37 HC) | Directional TB-score | **0.912** | 0.855–0.970 | 1.202 × 10<sup>−12</sup> | DeLong versus the 50-gene score, *p* = 0.0117; Holm (3 primary) *p* = 0.0234 |
| **Berry 393, unsigned** | GSE28623 (46 PTB + 37 HC) | Unsigned ssGSEA (*SEPT4*-corrected) | **0.828** | 0.740–0.916 | 7.56 × 10<sup>−8</sup> | 278 represented genes; DeLong versus the 50-gene score, *p* = 0.899 |
| **Berry 393, signed** | GSE28623 (46 PTB + 37 HC) | Signed ssGSEA (GSE19439 signs) | **0.910** | 0.844–0.975 | 1.95 × 10<sup>−12</sup> | Direction reconstructed, not the published Berry score; 204 up and 73 down represented; DeLong versus the 50-gene score, *p* = 0.0024, Holm *p* = 0.0071; signed versus unsigned *p* = 0.0037 |
| Berry 393, signed (sensitivity) | GSE28623 (46 PTB + 37 HC) | Signed ssGSEA (GSE19444 signs) | 0.911 | 0.846–0.975 | 1.61 × 10<sup>−12</sup> | Sign reconstruction check; DeLong versus the 50-gene score, *p* = 0.0018 |
| **50-gene score, Indonesia** | GSE56153 (18 PTB + 18 HC) | 50-gene ssGSEA (47/50 represented) | **0.941** | 0.868–1.000 | 4.595 × 10<sup>−7</sup> | External validation (wide CI) |

**Notes**
1. **DeLong method:** All AUC 95% confidence intervals and pairwise significance tests were computed using the DeLong asymptotic method.
2. **Sweeney 3-gene scoring:** Evaluated using the published formula (GBP5 + DUSP3)/2 − KLF2 on log2 expression values. This is a different scoring family from ssGSEA.
3. **Berry 393 scoring:** The published table has no per-gene direction column. The unsigned score uses ssGSEA across 278 represented genes after *SEPT4* correction (Excel date corruption 38231). Directions were reconstructed in GSE19439, where 225 genes were up-regulated and 86 down-regulated; 204 of the up-regulated and 73 of the down-regulated genes were represented in GSE28623 and entered the signed score. The signed score subtracts the enrichment of the down-regulated subset from that of the up-regulated subset; it is a reconstruction, not the published Berry score.
4. **Multiple testing:** The primary Holm family is the three comparator contrasts against the 50-gene score (Sweeney, unsigned Berry, signed Berry). Mean-z sensitivity rows are not in this family.
5. **Independence:** GSE107991 is a reanalysis of the Berry test set (GSE19444), Bloom is GSE42834, and both GSE42834 and GSE19444 (through GSE19491) were Sweeney discovery datasets. GSE28623 was not used to derive either signature, but it was a Sweeney validation cohort, so the head-to-head comparisons are not independent of signature derivation.
6. **4-gene classifier:** On the same 83 samples, the frozen 4-gene classifier (AUC 0.864) did not differ significantly from the Sweeney 3-gene score (paired DeLong *p* = 0.0557) or from the 50-gene score (*p* = 0.197).

Per-gene ROC analysis over the 47 testable genes gave a mean AUC of 0.747, with 15 genes (31.9%) at AUC ≥ 0.80 and 6 (12.8%) at AUC ≥ 0.85 (Table S4). The strongest single genes were *PSTPIP2* (0.900; 95% CI 0.831–0.968) and *LHFPL2* (0.883; 95% CI 0.805–0.961), neither a classical interferon-stimulated gene.

#### Direction concordance across three continents

Direction concordance was assessed in two further cohorts whose size precluded reliable AUC estimation. In GSE34608 (Germany; 8 PTB + 18 HC; Agilent GPL6480), 45 of 47 testable genes (95.7%) agreed with the discovery direction; the exceptions were *CACNA1E* and *SOCS1*. In GSE81746 (India; 4 PTB + 2 HC; Agilent GPL17077), 46 of 48 (95.8%) agreed; the exceptions were *GBP6* and *IDO1*. PCA for both cohorts is shown in Figure S7, and scale checks in Table S15. With GSE28623, these cohorts show directional consistency across three non-UK populations on three continents (Africa, Europe and Asia), on three Agilent platforms distinct from the Illumina discovery arrays. GSE81746, with 4 PTB and 2 HC, is descriptive only.

#### External validation (GSE56153, Indonesia)

The signature was also tested in GSE56153 (Ottenhoff et al. 2012; 18 PTB + 18 HC; Illumina GPL6883; Indonesia) [11]. Forty-seven of the 50 genes were represented on this platform after alias correction (absent: *ETV7*, *CMPK2*, *FCGR1CP*). ssGSEA gave an AUC of 0.941 (95% CI 0.868–1.000; Wilcoxon *p* = 4.595 × 10⁻⁷). The wide interval reflects the small sample (n = 36). The alias-corrected and old-symbol runs gave the same AUC and *p* value; the two added genes, *WARS1* and *IDO1*, shifted individual ssGSEA ranks by at most two positions (Spearman ρ = 0.998) without changing the case-control discrimination (Table S20).

### 3.6 Validation Stability and PTB versus LTBI Discrimination

We re-normalised all 108 GSE28623 samples together (46 PTB, 37 HC, 25 LTBI) and scored them by ssGSEA, to test stability under a different preprocessing choice and to compare active PTB with latent infection. The PTB-versus-HC AUC under 108-sample joint normalisation was 0.831 (95% CI 0.745– 0.918), a ΔAUC of +0.001 from the primary 83-sample value of 0.830 and inside the stability criterion of | ΔAUC| ≤ 0.05.

The signature separated active PTB from LTBI with an AUC of 0.848 (95% CI 0.756–0.940; Wilcoxon *p* = 3.238 × 10⁻⁷).

### 3.7 A Four-Gene Machine-Learning Classifier

#### Feature selection and model development

LASSO logistic regression was fitted to the 49 protein-coding consensus genes in the two microarray discovery cohorts (219 samples; 70 PTB, 149 HC). Four genes were chosen repeatedly: *GBP5* (5/5 folds), *BATF2* (5/5), *ANKRD22* (4/5) and *EPSTI1* (3/5). Forty-two of the 49 genes were never chosen, so most of the discriminative signal sits in a few genes.

Nested 5-fold cross-validation gave a mean AUC of 0.963 (SD 0.056; folds: 1.000, 1.000, 1.000, 0.876, 0.937). This measures feature sparsification and tuning inside a fixed 49-gene set, not the performance of an end-to-end discovery pipeline. A random forest trained on the 4 genes had an out-of-bag error rate of 3.02% on the SMOTE-augmented training data; because synthetic minority samples are included, this estimate is optimistic (Table 4B).

**TABLE 4A.** MCPcounter Differential Cell-Type Abundance Across Discovery Cohorts.

| **Cell Population** | **Direction in PTB** | **Concordance (significant studies)** | **GSE83456 adj. *p*** | **GSE83456 direction** | **Bloom adj. *p*** | **Bloom direction** | **GSE107991 adj. *p*** | **GSE107991 direction** |
|---|---|---|---|---|---|---|---|---|
| **T cells** | DOWN in PTB | **3/3** | 9.37 × 10<sup>−10</sup> | DOWN | 9.29 × 10<sup>−12</sup> | DOWN | 6.24 × 10<sup>−4</sup> | DOWN |
| **CD8 T cells** | DOWN in PTB | **3/3** | 1.10 × 10<sup>−3</sup> | DOWN | 4.22 × 10<sup>−4</sup> | DOWN | 3.68 × 10<sup>−2</sup> | DOWN |
| **Cytotoxic lymphocytes** | DOWN in PTB | **3/3** | 2.24 × 10<sup>−3</sup> | DOWN | 2.07 × 10<sup>−5</sup> | DOWN | 3.68 × 10<sup>−2</sup> | DOWN |
| **B lineage** | DOWN in PTB | **3/3** | 1.21 × 10<sup>−5</sup> | DOWN | 4.63 × 10<sup>−7</sup> | DOWN | 1.01 × 10<sup>−2</sup> | DOWN |
| **Monocytic lineage** | UP in PTB | **3/3** | 4.61 × 10<sup>−9</sup> | UP | 1.39 × 10<sup>−10</sup> | UP | 4.66 × 10<sup>−2</sup> | UP |
| **Endothelial cells** | UP in PTB | **3/3** | 1.39 × 10<sup>−2</sup> | UP | 3.39 × 10<sup>−3</sup> | UP | 2.33 × 10<sup>−2</sup> | UP |
| **Neutrophils** | UP in PTB | **2/3** | 2.58 × 10<sup>−7</sup> | UP | 7.19 × 10<sup>−8</sup> | UP | 8.24 × 10<sup>−2</sup> | (trend UP) |
| **NK cells** | DOWN in PTB | **2/3** | 4.91 × 10<sup>−3</sup> | DOWN | 3.63 × 10<sup>−4</sup> | DOWN | 0.340 | (trend DOWN) |
| **Myeloid dendritic cells** | Mixed / DOWN | **1/3** | 0.424 | DOWN | 0.984 | UP | 1.93 × 10<sup>−2</sup> | DOWN |
| **Fibroblasts** | Not significant | **0/3** | 0.424 | DOWN | 7.16 × 10<sup>−2</sup> | UP | 0.643 | DOWN |

**Notes**
1. **Deconvolution method:** Transcriptomic deconvolution performed using MCPcounter (10 immune and stromal cell populations).
2. **Hypothesis testing and multiple-testing correction:** Differences between PTB and HC evaluated by two-sided Wilcoxon rank-sum test with Benjamini-Hochberg (BH) false discovery rate control within each cohort. GSE83456 and Bloom were profiled on Illumina microarrays; GSE107991 by RNA-seq.
3. **Concordance criterion:** A cell population is classified as concordant if BH-adjusted *p* < 0.05 in at least two discovery cohorts with matching direction of effect. Eight of 10 cell populations (80%) meet this criterion.
4. **Immunological significance:** Myeloid populations expanded and lymphoid populations contracted, the pattern described in active TB [2].

**TABLE 4B.** Four-Gene Machine Learning Classifier Performance and Feature Importance.

| Evaluation Phase | Cohort / Analysis Context | Metric or Feature Parameter | Result / Value |
|---|---|---|---|
| **Feature Selection** | Discovery microarray pool (219 samples: 70 PTB, 149 HC) | LASSO L1 regularisation (family = "binomial", λ.1se) | Sparsification from 49 to 4 stable genes |
|  |  | ***BATF2*** (Selected in 5/5 outer CV folds, 100%) | Permutation Importance MDA = **21.61** |
|  |  | ***ANKRD22*** (Selected in 4/5 outer CV folds, 80%) | Permutation Importance MDA = **20.10** |
|  |  | ***GBP5*** (Selected in 5/5 outer CV folds, 100%) | Permutation Importance MDA = **20.00** |
|  |  | ***EPSTI1*** (Selected in 3/5 outer CV folds, 60%) | Permutation Importance MDA = **10.04** |
|  |  | Non-retained features | *FCGR1A* (2/5), *ETV7* (1/5), *GBP4* (1/5); 42 genes (0/5) |
| **Internal Validation** | 5-Fold Stratified Nested Cross-Validation | Nested CV Mean AUC ± SD | **0.963 ± 0.056** |
|  | (Outer 5-fold CV, Inner 10-fold λ tuning) | Per-fold outer AUCs | Fold 1: 1.000; Fold 2: 1.000; Fold 3: 1.000; Fold 4: 0.876; Fold 5: 0.937 |
|  | Final Random Forest fit (ntree = 500, mtry = 2) | Out-of-Bag (OOB) Error Rate | **3.02%** on SMOTE-augmented training data (optimistic) |
| **External Validation** | GSE28623 primary validation (46 PTB versus 37 HC; Agilent GPL4133) | Frozen 4-gene RF AUC (log2-then-z) | **0.864** (95% CI 0.785–0.944) |
|  | GSE28623 latent discrimination (46 PTB versus 25 LTBI) | Frozen 4-gene RF AUC (log2-then-z) | **0.867** (95% CI 0.784–0.949; Wilcoxon *p* = 4.164 × 10<sup>−8</sup>) |
| **Imbalance Sensitivity** | Inverse Frequency Weighting (IFW) + SMOTE | Primary Dual Imbalance Correction | AUC = 0.963 ± 0.056 |
|  | IFW-only (no SMOTE) | Ablation Variant 1 | AUC = 0.962 ± 0.056 (ΔAUC = −0.001) |
|  | SMOTE-only (no IFW) | Ablation Variant 2 | AUC = 0.963 ± 0.056 (ΔAUC = 0.000) |

**Notes**
1. **Model architecture:** Feature sparsification by LASSO penalised logistic regression followed by classification with a random forest (ntree = 500, mtry = 2).
2. **Feature collinearity:** Network analysis identified 10 hub genes co-regulated in the interferon module (*STAT1*, *IFI35* and others). LASSO selected only *EPSTI1* among them. The other three features were less correlated with the hub set: *BATF2* and *GBP5* are interferon-module genes outside the 10 hubs, and *ANKRD22* is in the other module. We describe the set as low-correlation or non-redundant for classification.
3. **External validation:** The frozen 4-gene RF classifier was evaluated on GSE28623 without retraining or tuning. Because the external matrix is on a linear intensity scale while the training matrices are log2-scale, log2 transformation preceded per-gene z-scoring.
4. **Out-of-bag error:** The 3.02% OOB error was computed on the 298-sample SMOTE-balanced training set. Because synthetic minority samples are included, it is an optimistic internal estimate, not an external validation estimate.

#### External validation

The frozen 4-gene random forest was applied to the held-out GSE28623 cohort with per-gene z-score normalisation. Because the external matrix is on a linear intensity scale and the training matrices are on log2 scale, we log2-transformed before z-scoring. The classifier reached an AUC of 0.864 (95% CI 0.785– 0.944; Table S13). That figure is not directly comparable with the 50-gene ssGSEA AUC of 0.830: the forest uses four LASSO-selected genes, whereas ssGSEA measures enrichment of the whole 50-gene module.

#### Feature importance and hub gene comparison

By permutation importance (mean decrease in accuracy) the order was *BATF2* (21.61), *ANKRD22* (20.10), *GBP5* (20.00) and *EPSTI1* (10.04) (Table S6). *EPSTI1* was the only gene in both the top-10 CytoHubba set and the final 4-gene model (1/10 overlap). The remaining 9 hubs (*STAT1*, *IFI35*, *GBP1*, *IFIT3*, *RSAD2*, *ISG15*, *XAF1*, *SAMD9L* and *OAS3*) were selected in 0 of the 5 outer folds, because LASSO discards collinear hub genes in favour of less correlated genes, including interferon-module genes outside the hub set.

#### Frozen RF validation against LTBI

The frozen 4-gene random forest, with the same log2-then-z preprocessing, separated PTB (n = 46) from LTBI (n = 25) in GSE28623 without retraining, SMOTE or tuning, at an AUC of 0.867 (95% CI 0.784– 0.949; Wilcoxon *p* = 4.16 × 10⁻⁸).

#### Imbalance correction sensitivity

IFW+SMOTE (primary; AUC 0.963), IFW alone (0.962) and SMOTE alone (0.963) differed by at most 0.001, so the result did not depend on the choice of imbalance correction (Table S7).

### 3.8 Comparison with Published TB Signatures

We compared the 50-gene signature with two established blood signatures on the same 83 GSE28623 samples, using paired DeLong tests (Table 3). Each signature kept its own scoring family: ssGSEA for the 50-gene score and both Berry scores, and the published directional formula for the Sweeney 3-gene score.

The Sweeney 3-gene score (*GBP5*, *DUSP3*, *KLF2*; directional TB score) reached an AUC of 0.912 (95% CI 0.855–0.970; Wilcoxon *p* = 1.202 × 10⁻¹²), above the 50-gene ssGSEA score (paired DeLong *p* = 0.0117; Holm across the three primary comparisons *p* = 0.0234). The frozen 4-gene classifier reached 0.864 on the same samples and did not differ significantly from the Sweeney score (paired DeLong *p* = 0.0557) or from the 50-gene score (*p* = 0.197).

The Berry 393-gene set was scored two ways because the published table has no direction column (Section 2.11). Scored without direction over all 278 represented genes, it gave an AUC of 0.828 (95% CI 0.740– 0.916; Wilcoxon *p* = 7.56 × 10⁻⁸), indistinguishable from the 50-gene score (DeLong *p* = 0.899). The signed score, which subtracts the enrichment of the 73 genes represented in GSE28623 and reconstructed as down-regulated from that of the 204 up-regulated genes, gave an AUC of 0.910 (95% CI 0.844–0.975; Wilcoxon *p* = 1.95 × 10⁻¹²) and exceeded both the unsigned Berry score (DeLong *p* = 0.0037) and the 50-gene score (DeLong *p* = 0.0024; Holm *p* = 0.0071; Table S14). Signs reconstructed separately in GSE19444 gave a nearly identical score (0.911; 95% CI 0.846–0.975). The signed Berry score is a reconstruction, not Berry’s published per-gene score; the two versions belong side by side.

Both the Sweeney 3-gene score and the signed Berry score ranked above the 50-gene score, which could not be distinguished from the unsigned Berry score. The nominal Sweeney 3-gene versus unsigned Berry 393-gene comparison gave *p* = 0.027, unadjusted (Table S17). Mean-z versions of the Berry score gave 0.914 (signed, GSE19439 signs) and 0.815 (unsigned); these are sensitivity analyses and were left out of the primary correction.

Four caveats apply. The comparisons rest on one validation cohort, and scoring methods differ (ssGSEA against a directional formula), as do gene-set sizes (50, 393 and 3 genes). Finally, the signatures were not derived from separate cohorts: GSE107991 reanalyses the Berry test set (GSE19444), Bloom is GSE42834, and both GSE42834 and GSE19444 (through GSE19491) were Sweeney discovery datasets [2, 3]. GSE28623 was not used to derive either signature, so the AUCs remain external to it, but the head-to-head is not a comparison of independently derived signatures.

### 3.9 Immune Cell Deconvolution

#### MCPcounter

MCPcounter identified 8 of 10 cell populations as altered in PTB (BH-adjusted *p* < 0.05) in at least 2 of the 3 discovery cohorts (Figure 7, Table 4A; Table S8). Six populations were concordant in all three: T cells, CD8 T cells, cytotoxic lymphocytes and B lineage, all lower in PTB, and monocytic lineage and endothelial cells, both higher. Neutrophils (higher) and NK cells (lower) agreed in 2 of 3. Figure S8 shows per-cohort boxplots for Bloom and GSE107991, complementing the GSE83456 panels in Figure 7B.

**Figure 7.**
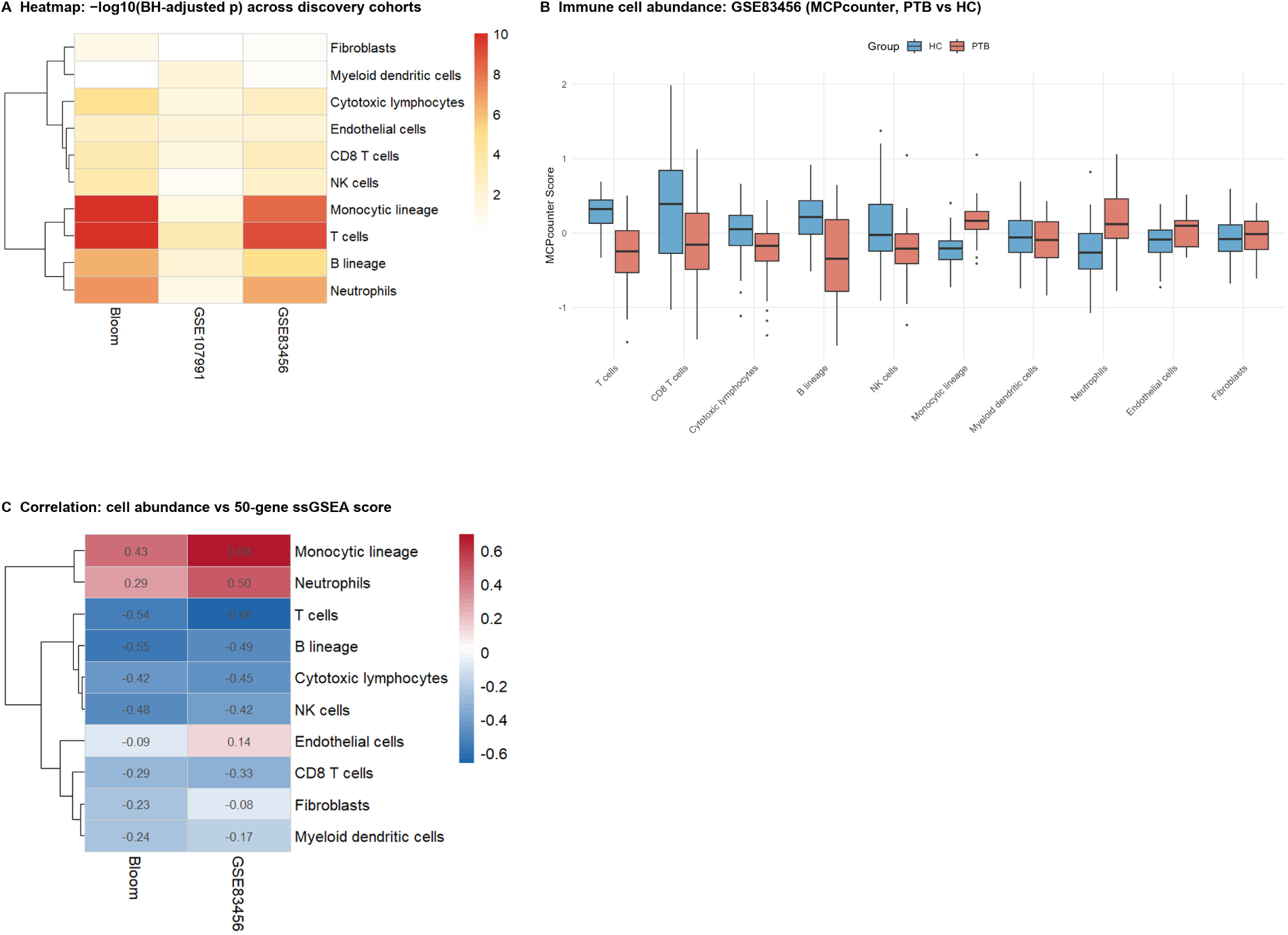
Immune cell deconvolution. MCPcounter scores were compared across the discovery cohorts. (A) Heatmap of −log10 BH-adjusted p-values for differential cell-type abundance (PTB versus HC) across the three cohorts. Six cell populations were concordantly altered across all three datasets. (B) Representative boxplots of MCPcounter scores in GSE83456, showing monocytic lineage expansion and T cell suppression in PTB. (C) Correlation between MCPcounter cell-type scores and the 50-gene ssGSEA enrichment score. Monocytic lineage showed the strongest positive correlation (Spearman ρ = +0.693; BH p < 2.2 × 10⁻¹⁶), while T cells showed a strong negative correlation (ρ = −0.659).

#### CIBERSORTx

CIBERSORTx with the LM22 reference (22 immune cell subtypes) found 5 cell types altered in at least two cohorts (Table S9): monocytes up (3/3), M1 macrophages up (2/3), M2 macrophages up (2/3), activated dendritic cells up (2/3) and T follicular helper cells down (2/3). QC pass rates were 42.7% in GSE83456, 57.8% in Bloom and 100% in GSE107991. The low microarray rates reflect platform limits against the LM22 reference and leave CIBERSORTx comparisons underpowered.

#### Cross-method concordance

MCPcounter showed myeloid expansion (monocytes and neutrophils) and lymphoid suppression (T cells, CD8 T cells, B cells and NK cells), the pattern described in active TB [2, 5]. CIBERSORTx recovered the myeloid expansion (monocytes, M1 and M2 macrophages and activated dendritic cells) and lower T follicular helper cells, but it did not confirm the broader lymphoid suppression. The two methods therefore agree on the myeloid signal and diverge on the lymphoid one. Within the myeloid compartment, both M1 and M2 macrophage scores were raised, but the LM22 estimates are relative and the microarray pass rates were low, so we do not interpret this as evidence of polarisation.

#### Mechanistic link to the consensus signature

MCPcounter abundances tracked the 50-gene ssGSEA score in the discovery cohorts. In GSE83456, monocytic lineage correlated positively (Spearman ρ = +0.693; BH *p* < 2.2 × 10⁻¹⁶) and T cells negatively (ρ = −0.659; BH *p* < 2.2 × 10⁻¹⁶); neutrophils correlated more weakly (ρ = +0.500; BH *p* = 3.97 × 10⁻⁷). The score therefore follows monocyte and T-cell abundance in part. Correlation with deconvolution does not establish mechanism (Table S10).

### 3.10 Summary of Validation Performance

Table 3 brings together the validation estimates from Sections 3.5 to 3.8. The 50-gene score, computed over the 47 genes represented in each cohort, gave AUCs of 0.830 in GSE28623 and 0.941 in GSE56153; the frozen 4-gene classifier gave 0.864 for PTB versus HC and 0.867 for PTB versus LTBI; the Sweeney 3-gene score gave 0.912 in GSE28623. Paired DeLong comparisons are given in the table notes.

## 4. Discussion

### 4.1 Summary of Principal Findings

Meta-analysis of three discovery datasets on two platforms, Illumina microarray and RNA-seq, produced a 50-gene blood transcriptional signature for active pulmonary tuberculosis. One dataset, GSE107991, is a reanalysis of the London test set used to validate the Berry 393-gene signature, so the three are discovery sets, not independent cohorts. Performance replicated outside the UK, with AUCs from 0.830 in The Gambia to 0.941 in Indonesia. ssGSEA scores gave an AUC of 0.830 (95% CI 0.743–0.917) in The Gambia (GSE28623) and 0.941 (95% CI 0.868–1.000) in Indonesia (GSE56153), and direction concordance was 95.7% in the smaller German cohort and 95.8% in the Indian cohort. A 4-gene random forest (*BATF2*, *ANKRD22*, *GBP5*, *EPSTI1*), built by sparsifying the consensus candidates, reached an internal nested cross-validation AUC of 0.963 within the fixed 49-gene set and an external AUC of 0.864 (95% CI 0.785–0.944). Without retraining it separated PTB from LTBI at an AUC of 0.867 (95% CI 0.784–0.949). MCPcounter showed myeloid expansion and lymphoid suppression, whereas CIBERSORTx recovered the myeloid expansion but not the broader lymphoid suppression.

### 4.2 Relationship to Established TB Signatures

The 50-gene signature is embedded in the classical TB interferon response: 42 of the 50 genes are in the Berry et al. (2010) 393-transcript list. This study is best read as a reproducibility and benchmarking analysis, not as a claim that the 50-gene score is the best available test. All 10 CytoHubba hubs are interferon-stimulated genes described in the Berry et al. (2010) 393-transcript signature [2], so the dominant signal is a type I/II interferon response, the most reproducible feature of the host blood response to active TB [2, 5].

In GSE28623 the Sweeney et al. (2016) 3-gene score [3] reached an AUC of 0.912, above the 50-gene ssGSEA score of 0.830 (paired DeLong *p* = 0.0117; Holm across the three primary comparisons *p* = 0.0234). The frozen 4-gene classifier (AUC 0.864) did not differ significantly from the Sweeney score (paired DeLong *p* = 0.0557). The Berry 393-gene set has no single comparator value, because the published table gives no per-gene direction. Scored without direction over the 278 represented genes, it gave an AUC of 0.828, indistinguishable from the 50-gene score (DeLong *p* = 0.899). Scored with directions reconstructed from the Berry training set (GSE19439; 225 up and 86 down there, of which 204 and 73 were represented in GSE28623), it gave 0.910 and exceeded the 50-gene score (DeLong *p* = 0.0024; Holm *p* = 0.0071). The signed score is a reconstruction, not Berry’s published score, so both versions are reported.

Sweeney’s score is a directional formula built for binary classification from three genes that point in opposite directions; ssGSEA measures coordinated enrichment of a whole module. Gene-set size (3 against 50 and 393) further complicates a direct comparison. In this one cohort, the signed Berry score’s advantage over the unsigned score suggests that direction-aware scoring captures the module better, and the Sweeney result suggests that a small, directionally optimised score can classify better than module-level enrichment [3]. Neither comparison is independent. GSE107991 reanalyses the Berry test set, and the Bloom dataset (GSE42834) and GSE107991 (through GSE19444 within GSE19491) overlap with the datasets Sweeney used for discovery [2, 3]. GSE28623 was not used to derive either signature, so the performance estimates stand, but the two signatures are not products of separate discovery programmes. We therefore present these comparisons as benchmarks, not as evidence that the new scores are better.

### 4.3 Candidate Biomarkers Outside the Interferon Axis

Two genes outside the interferon axis are worth a closer look, but both were selected as the top two single-gene AUCs among the 47 testable genes, so the observations are exploratory. *PSTPIP2* (proline-serine-threonine phosphatase-interacting protein 2) gave the highest single-gene AUC in the Gambian cohort (0.900). *PSTPIP2* has been linked to NLRP3 inflammasome regulation [37], and an AUC at that level is compatible with a detectable secondary signal, although selection makes the estimate optimistic. *Mycobacterium tuberculosis* ESAT-6 triggers NLRP3 inflammasome activation and IL-1β secretion in host cells [38], so the rise in *PSTPIP2* could be a regulatory counter-response, but this is a hypothesis. The signal appeared in a geographically distinct cohort, which makes a UK-specific artefact less likely, but its mechanism remains untested.

*LHFPL2* (lipoma HMGIC fusion partner-like 2), a tetraspan transmembrane protein, gave the second-highest AUC (0.883). It is an isolate in the PPI network, with no STRING interactions at medium confidence, and has no well-established role in TB immunology or interferon signalling. Why it is consistently upregulated across platforms and populations is unexplained. We put it forward as an exploratory candidate that needs functional characterisation.

### 4.4 Network Topology versus Machine Learning: Complementary Perspectives

The network hubs and the machine-learning features overlapped little. *EPSTI1* was the only gene in both the CytoHubba MCC top 10 and the final 4-gene LASSO model (1/10 overlap). CytoHubba and LASSO optimise different properties. CytoHubba MCC rewards topological centrality inside the co-expressed interferon module and therefore favours highly connected proteins such as *STAT1* and *ISG15*. LASSO with L1 regularisation tends to select one representative from each group of correlated features and discard the rest. Hub genes can therefore be central to the biology yet redundant for classification, and LASSO resolves that redundancy by keeping one representative per cluster. *EPSTI1* is the one gene that is both central in the network and non-redundant for classification. These data cannot settle whether that makes it the best single marker.

### 4.5 Immune Deconvolution and Mechanistic Interpretation

MCPcounter showed myeloid expansion (monocytes and neutrophils) with lymphoid suppression (T cells, CD8 T cells, B cells and NK cells), a pattern consistent with the myeloid expansion and lymphoid suppression described in active TB [2, 5]. CIBERSORTx recovered the myeloid signal but not the broader lymphoid suppression. The divergence may be methodological as much as biological: MCPcounter estimates relative abundance from lineage markers, whereas CIBERSORTx fits a 22-cell reference matrix, and the low LM22 pass rates in the microarray cohorts limit the precision of the CIBERSORTx estimates. The myeloid expansion is the more robust finding because both methods recover it and MCPcounter detects it in all three discovery cohorts. The correlation between monocytic lineage and the 50-gene score (Spearman ρ = +0.693) shows that the score partly follows monocyte abundance, not interferon activity alone. The endothelial signal should be interpreted cautiously: in whole blood it may reflect platelet or vascular contamination rather than pulmonary endothelium.

CIBERSORTx added resolution within the myeloid compartment: both M1 and M2 macrophage scores were raised. Because the LM22 estimates are relative and the microarray pass rates were low, we do not interpret this as evidence of polarisation. *M. tuberculosis* can push macrophage polarisation toward an M2-like phenotype in experimental systems [39], but whole-blood deconvolution cannot establish that this occurs in the patients studied here.

### 4.6 Design of the Discovery and Validation Steps

Gene selection did not rely on pooling alone. Every gene in the 50-gene signature reached FDR < 0.05 independently in all three datasets before meta-analysis set its effect size. The |logFC| > 1.5 cutoff retained the top 50 of the 1,959 replicated genes (97.4th percentile), which gives it an empirical basis instead of an arbitrary one. No gene selection or tuning touched the validation data, so the external cohorts provide an independent test of generalisability for the 50-gene signature. Network analysis (STRING, CytoHubba, MCODE), machine learning (LASSO, random forest, permutation importance) and deconvolution (MCPcounter, CIBERSORTx) ran in one workflow, which connects discovery, mechanism and classifier development.

### 4.7 Limitations

All three discovery datasets are from the UK, which may limit generalisability. External evaluation in four cohorts from The Gambia, Germany, India and Indonesia (two with AUC estimation, two with direction concordance only) partly addresses this. GSE107991 is a reanalysis of the Berry test set, and two of the discovery datasets overlap Sweeney’s discovery data (Section 2.11). These overlaps do not change the within-study replication criterion, but they make both head-to-head comparisons non-independent. The studies share no subject-level identifier, so patient overlap between GSE83456, Bloom and the Berry London cohorts cannot be excluded; this is an inherent limitation of secondary analysis of public data.

The study uses public data only and adds no local qPCR cohort. Sarcoidosis samples in GSE34608 and other-disease samples in GSE42834 were excluded rather than analysed, so no non-TB disease comparison group remained; the scores have therefore not been tested against pneumonia, sarcoidosis or other mycobacterial infections, which is the main threat to clinical validity. We also did not perform fixed-threshold sensitivity/specificity analyses or the meta-analysis sensitivity analyses (unshrunk effect sizes and standard errors, fixed-effect, HKSJ, leave-one-out and alternative cutoffs) planned for the journal revision. These are design and analysis limits, not evidence that the current estimates are definitive.

Three studies limit how well between-study variance can be estimated. We did not use the HKSJ correction for primary inference: with k = 3 the t-based intervals are wide and may be conservative under substantial heterogeneity [17, 18]. The safeguard was the replication criterion, which requires all 50 genes at FDR < 0.05 in every study and does not depend on the pooling method. Twelve of the 50 core genes (24%) had I² ≥ 75%, although I² is unstable with k = 3 [19]. We pooled microarray and RNA-seq log2 fold-changes directly, without converting to Hedges’ g. Both platforms give log2 fold-changes and the random-effects τ² absorbs residual variance [16], but formal standardisation would be better in future work.

CIBERSORTx pass rates in the microarray cohorts were low (42.7% and 57.8%), reflecting platform-specific limitations against the LM22 reference; statistical power for CIBERSORTx-specific findings is therefore reduced, and CIBERSORTx did not confirm the lymphoid suppression seen with MCPcounter, so that component rests on one method. HIV status was not available for GSE56153 and may confound the high AUC in that cohort.

*CACNA1E* remains unexplained. It is a PPI isolate with no established TB mechanism, shows directional discordance in the German cohort and had a 0% LASSO selection frequency; we kept it for completeness. The 49 candidate genes given to machine learning came from the discovery meta-analysis, so the nested cross-validation AUC (0.963; SD 0.056) measures feature sparsification inside that fixed set, not an end-to-end discovery pipeline. The external AUC in GSE28623 (0.864) is the appropriate benchmark for the 4-gene classifier.

### 4.8 Clinical Implications and Future Directions

Neither the 50-gene signature nor the 4-gene classifier is ready for clinical use. Clinical translation would need prospective validation in endemic populations, testing against differential diagnoses (pneumonia, sarcoidosis, other mycobacterial infections), assessment across HIV co-infection status, and adaptation to a point-of-care platform. For now the signature is best read as a reproducible molecular phenotype of the host response to active PTB, useful for prioritising candidates and for pathway work.

Future work should test the signature in larger, geographically diverse prospective cohorts with confirmed diagnoses, measure performance in HIV co-infection, and establish what *PSTPIP2* and *LHFPL2* do in TB immunopathology. The 4-gene classifier (*BATF2*, *ANKRD22*, *GBP5*, *EPSTI1*) could be developed into an RT-qPCR assay, but it first needs validation outside this computational framework.

## Data Availability Statement

Supplementary figures S1–S9 accompany this preprint as a separate Supplementary Material file, which also indexes supplementary tables S1–S20. The tables themselves are archived with the analysis code and data on Zenodo (concept DOI: 10.5281/zenodo.23808440; version 1.3.0). The four tables affected by the HGNC symbol correction (S04, S15b, S15c and S17) and the two v1.3 verification tables (S19 and S20) are reproduced in Appendix A of the supplement; the earlier v1.0.0 version remains available and contains the pre-correction versions of the four tables.

All transcriptomic datasets analysed in this study are publicly available in the NCBI Gene Expression Omnibus (GEO) repository under accession numbers GSE83456 (https://www.ncbi.nlm.nih.gov/geo/query/acc.cgi?acc=GSE83456), GSE42834 (comprising sub-series GSE42826 and GSE42830; https://www.ncbi.nlm.nih.gov/geo/query/acc.cgi?acc=GSE42834), GSE107991 (https://www.ncbi.nlm.nih.gov/geo/query/acc.cgi?acc=GSE107991), GSE28623 (https://www.ncbi.nlm.nih.gov/geo/query/acc.cgi?acc=GSE28623), GSE34608 (https://www.ncbi.nlm.nih.gov/geo/query/acc.cgi?acc=GSE34608), GSE81746 (https://www.ncbi.nlm.nih.gov/geo/query/acc.cgi?acc=GSE81746), and GSE56153 (https://www.ncbi.nlm.nih.gov/geo/query/acc.cgi?acc=GSE56153). Processed expression matrices, phenotype tables, consensus gene lists and frozen result tables are archived on Zenodo (concept DOI: 10.5281/zenodo.23808440; version 1.3.0).

## Code Availability Statement

All analysis code is archived on Zenodo (concept DOI: 10.5281/zenodo.23808440; version 1.3.0). The deposit contains the scripts for differential expression, random-effects meta-analysis, ssGSEA scoring, classifier training, enrichment analysis and immune deconvolution, together with a SHA-256 manifest of every deposited file and a claims ledger that records, for each quantitative result, its value, evidence path, locator and verification status. A fail-closed script, audit_check.R, verifies that every ledger entry has an extant evidence file, that entries marked as reproduced carry the command that regenerates them, and that all frozen inputs match their recorded SHA-256 checksums; it exits non-zero on any failure.

## Ethics Statement

This study is a secondary analysis of publicly available, de-identified human transcriptomic data deposited in the NCBI GEO repository. Ethical approval and participant informed consent were obtained by the principal investigators of the primary studies cited herein.

## Author Contributions

**Usama Hassan:** Conceptualisation, Data curation, Formal analysis, Investigation, Methodology, Project administration, Software, Validation, Visualisation, Writing – original draft. **Farah Tahir:** Investigation, Writing – review & editing. **Armeela Bibi:** Data curation, Formal analysis, Writing – review & editing. **Warda Jamal:** Data curation, Investigation, Writing – review & editing. **Tooba Kanwal:** Investigation, Writing – review & editing.

All authors have read and approved the preprint and the Zenodo deposit and agree to release them under CC BY 4.0.

## Use of Artificial Intelligence

The authors used an AI language model to assist with manuscript editing, consistency checking, and code and figure verification. The model did not generate data, analyses or references. All scientific content, numerical results, interpretations and final wording were reviewed and approved by all authors, who take responsibility for the work.

## Acknowledgments

We thank the primary investigators and study participants who made the transcriptomic datasets publicly accessible through the NCBI Gene Expression Omnibus.

