## Supplementary Material for "Multi-Cohort Blood Transcriptomic Meta-Analysis of Active Pulmonary Tuberculosis: Reproducibility of a 50-Gene Interferon Signature and Benchmarking of a 4-Gene Classifier"

Version 1.3 · 12 November 2026

[1]Department of Medical Lab Technology, University of Haripur, Haripur, Khyber Pakhtunkhwa, Pakistan; [2]Faculty of Rehabilitation and Allied Health Sciences, Riphah International University, Al-Mizan Campus, Rawalpindi, Punjab, Pakistan

This supplement contains Figures S1–S9, an index of Supplementary Tables S1–S20, and Appendix A, which reproduces the four tables affected by the v1.2 HGNC symbol correction (S04, S15b, S15c, S17), together with the v1.3 module annotation (S19) and the GSE56153 alias comparison (S20). The remaining table files are archived, with checksums, in the Zenodo deposit (concept DOI: 10.5281/zenodo.23808440; version 1.3.0). The earlier v1.0.0 version remains available and carries the pre-correction versions of the four tables. Version 1.3 retains the corrected mapping introduced in v1.2 (WARS → WARS1; INDO → IDO1).

---

#### Contents

1. **Figure S1.** Quality control, GSE83456
2. **Figure S2.** Quality control, Bloom
3. **Figure S3.** Quality control, GSE107991
4. **Figure S4.** Boxplots before and after normalisation
5. **Figure S5.** KEGG GSEA, three discovery datasets
6. **Figure S6.** Discovery ssGSEA boxplots (apparent)
7. **Figure S7.** PCA for direction-only cohorts
8. **Figure S8.** MCPcounter boxplots, additional cohorts
9. **Figure S9.** Full ORA GO dotplot

#### Supplementary figures

##### Figure S1

Quality control, GSE83456

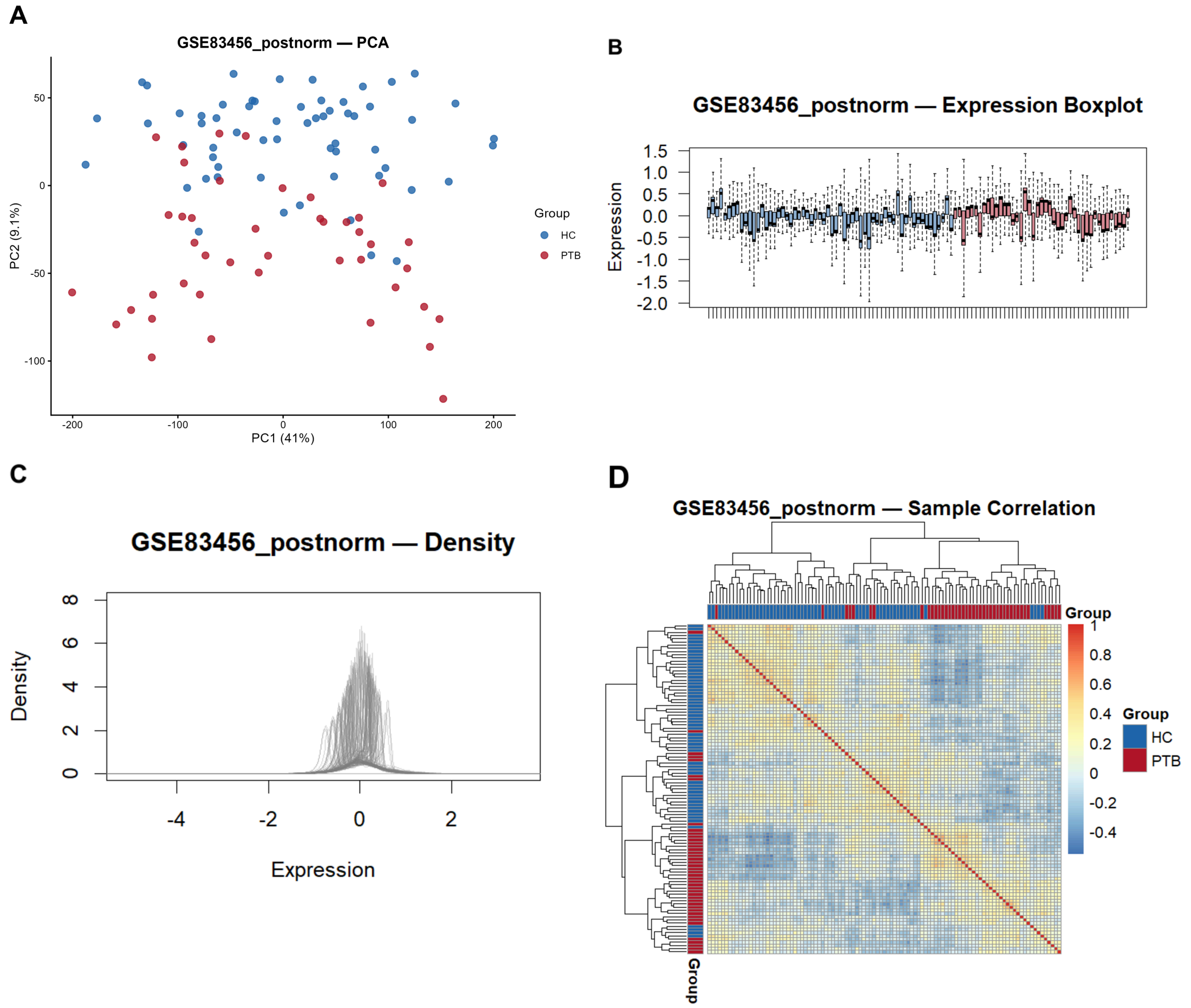


**Figure S1.** Post-normalisation quality control for GSE83456 (103 samples; 43 PTB, 60 HC). (A) PCA coloured by disease group. (B) Per-sample expression boxplots. (C) Expression density distributions. (D) Hierarchical clustering and sample–sample correlation heatmap with disease annotation.

#### Figure S2

Quality control, Bloom

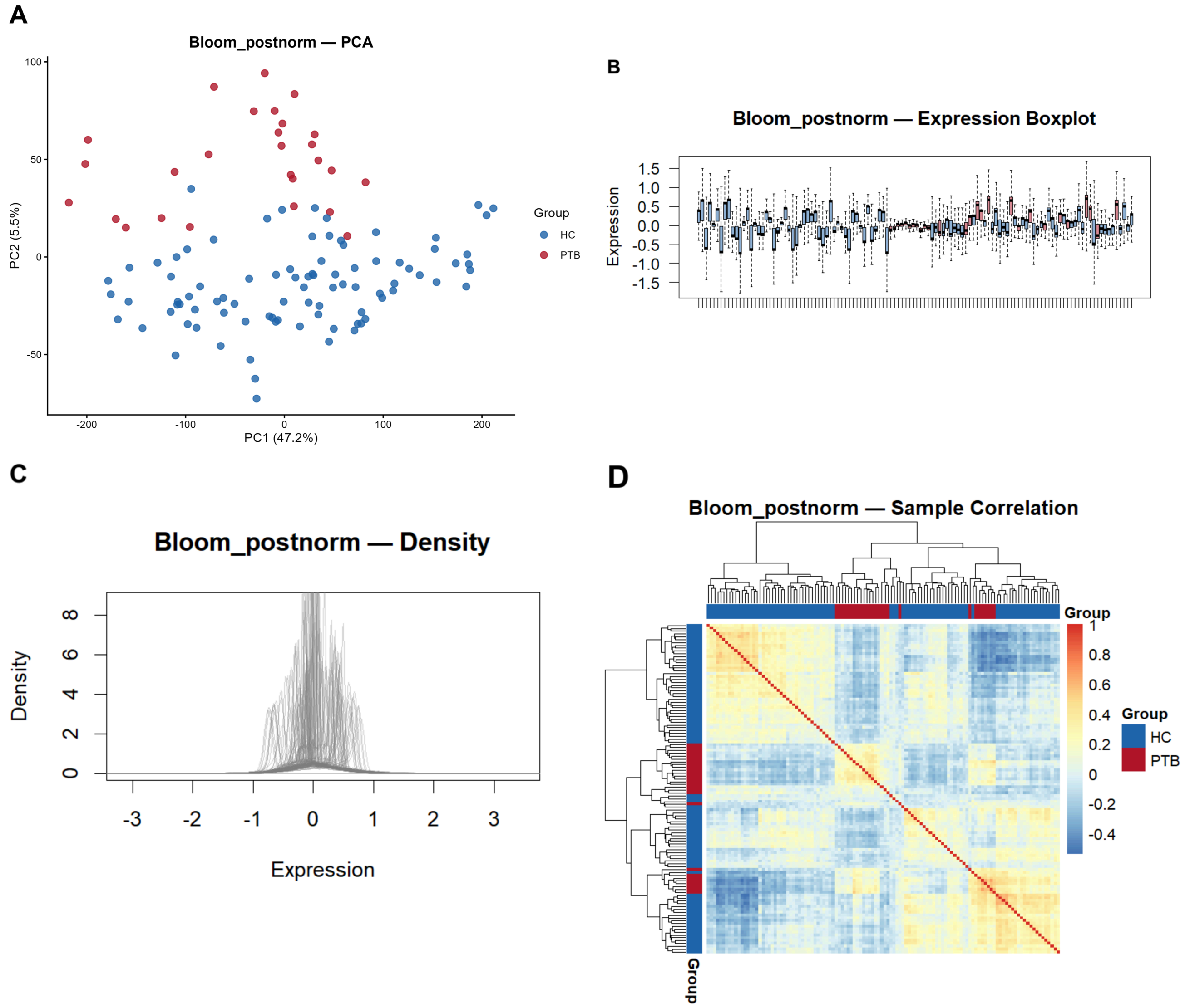


**Figure S2.** Post-normalisation quality control for the merged Bloom dataset (116 samples; 27 PTB, 89 HC). Panels as in Figure S1. Batch structure for the two sub-series is shown in main Figure 1D.

#### Figure S3

Quality control, GSE107991

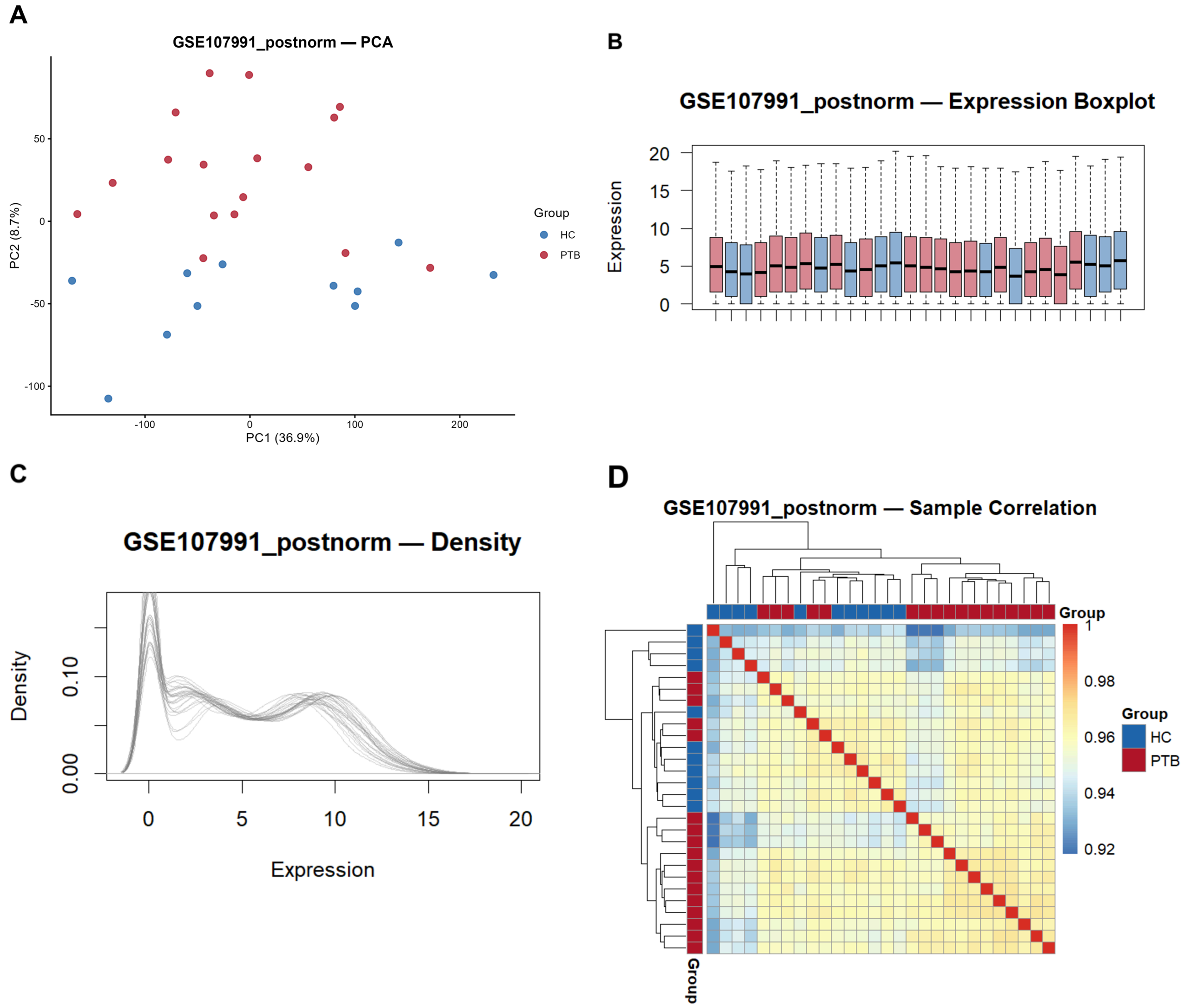


**Figure S3.** Post-normalisation quality control for GSE107991 (28 samples; 17 PTB, 11 HC). Panels as in Figure S1.

### Figure S4

Boxplots before and after normalisation

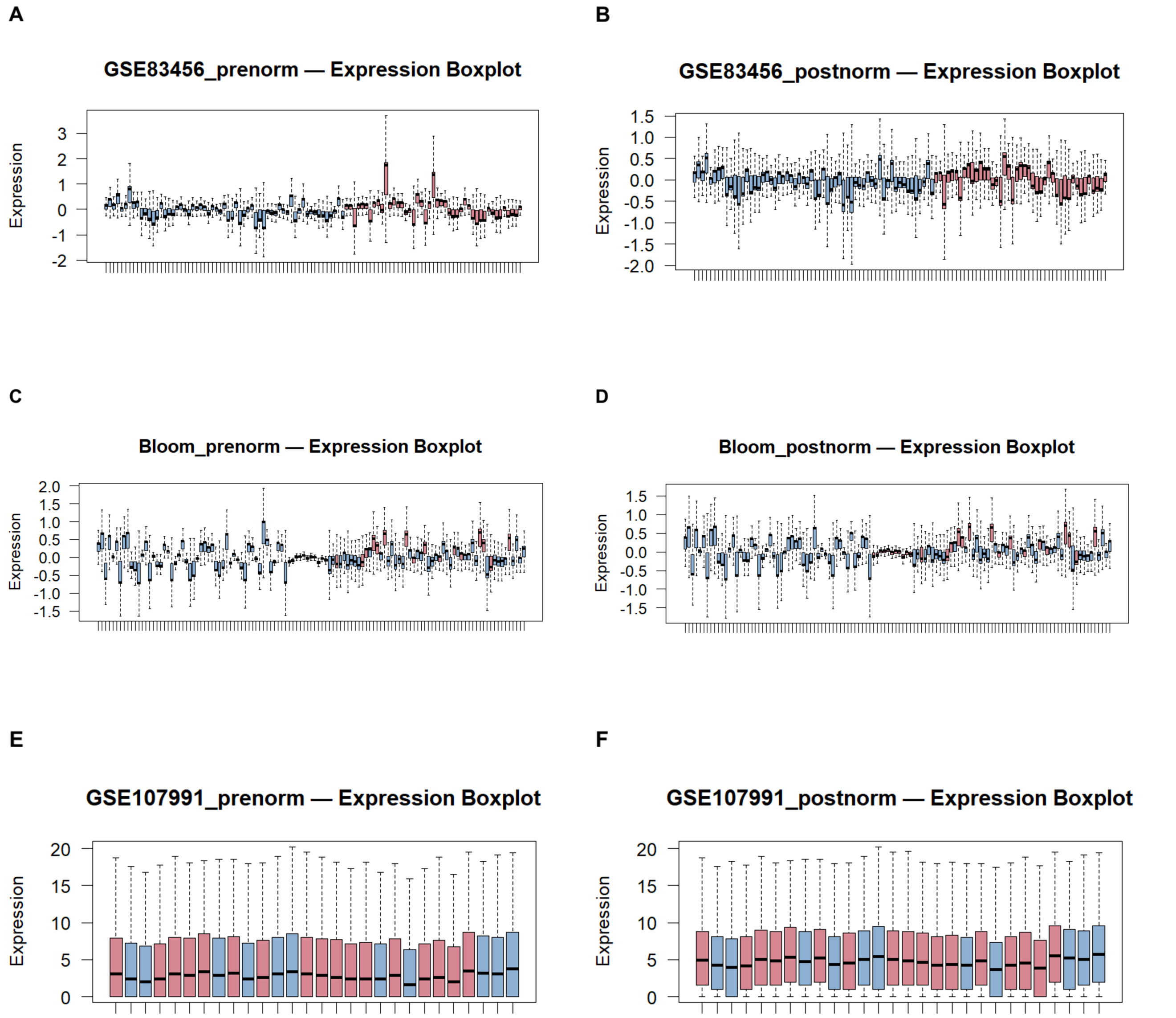


**Figure S4.** Per-sample expression boxplots before and after normalisation for GSE83456 (A, B), Bloom (C, D), and GSE107991 (E, F). GSE107991 is RNA-seq and retains a wider expression range after normalisation.

### Figure S5

KEGG GSEA, three discovery datasets

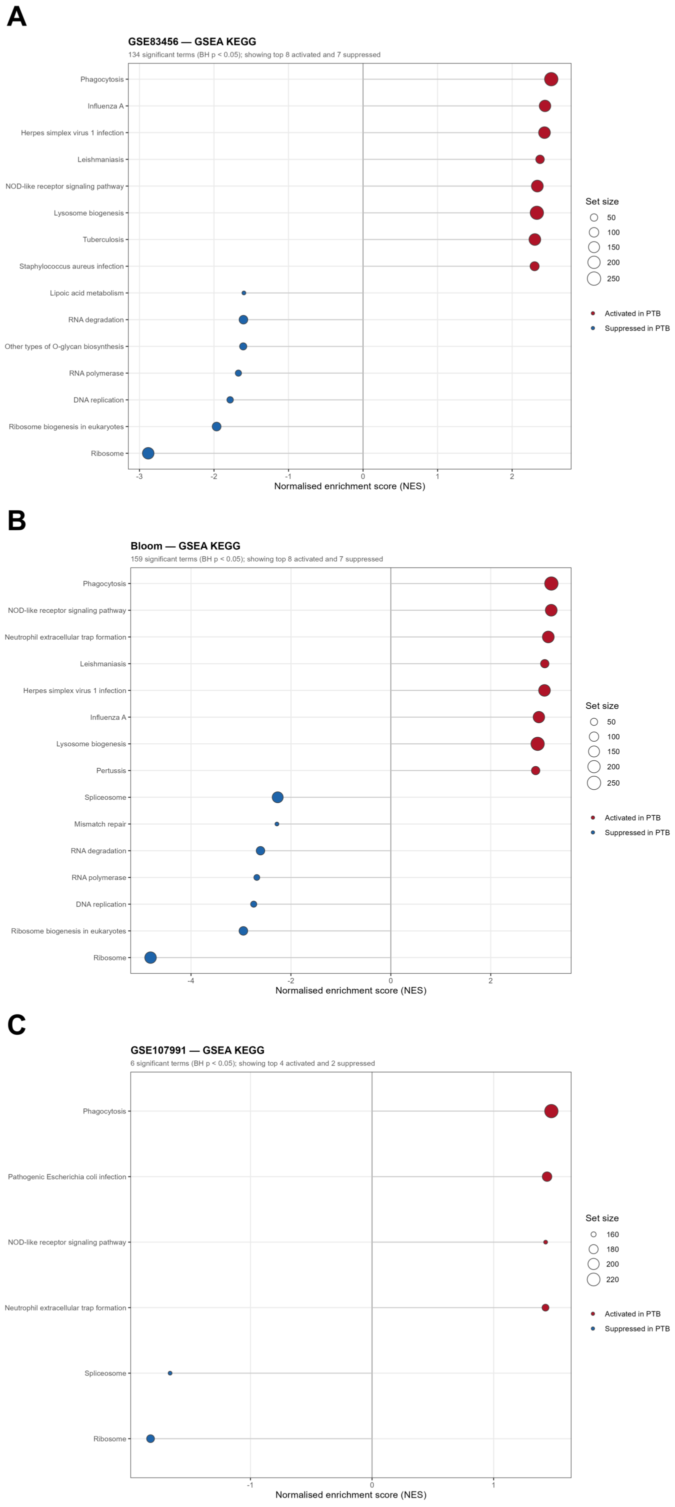


**Figure S5.** KEGG pathway GSEA in each discovery dataset, drawn with the same convention as Figure 3. (A) GSE83456 (134 significant pathways). (B) Bloom (159). (C) GSE107991 (6). Each panel shows the eight most strongly activated and seven most strongly suppressed pathways at BH FDR < 0.05. Ribosome and ribosome-biogenesis pathways are suppressed, while phagocytosis, lysosome, NOD-like receptor, and TNF signalling are activated.

#### Figure S6

Discovery ssGSEA boxplots (apparent)

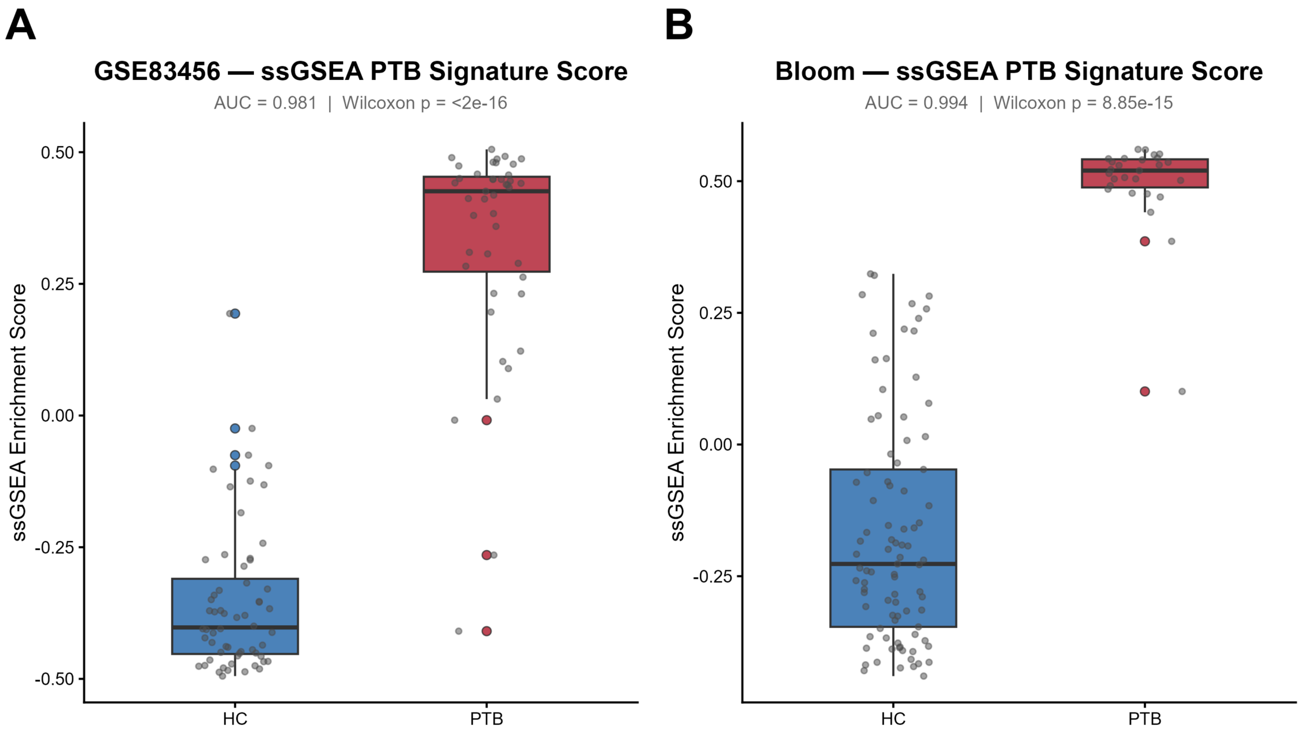


**Figure S6.** Apparent (resubstitution) ssGSEA enrichment scores for the 50-gene signature in the two discovery microarray cohorts: (A) GSE83456 and (B) Bloom. These are within-discovery values and are not external validation estimates.

#### Figure S7

PCA for direction-only cohorts

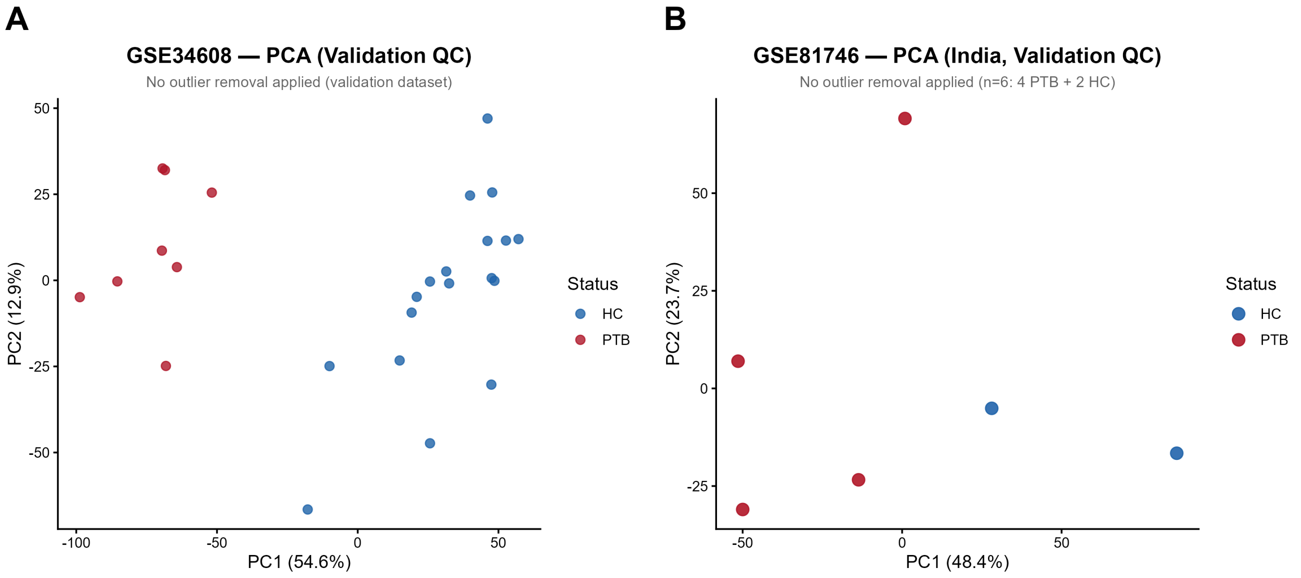


**Figure S7.** PCA of (A) GSE34608 (26 samples; 8 PTB, 18 HC) and (B) GSE81746 (6 samples; 4 PTB, 2 HC). Both cohorts were used for direction concordance only; no AUC was computed.

### Figure S8

MCPcounter boxplots, additional cohorts

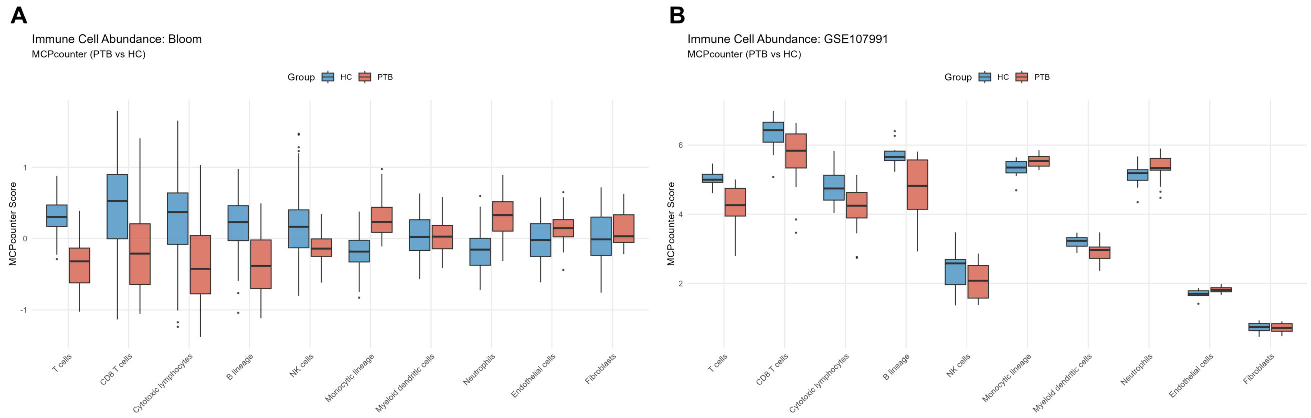


**Figure S8.** MCPcounter cell-type scores in (A) Bloom and (B) GSE107991, complementing the GSE83456 boxplots in main Figure 7B.

#### Figure S9

Full ORA GO dotplot

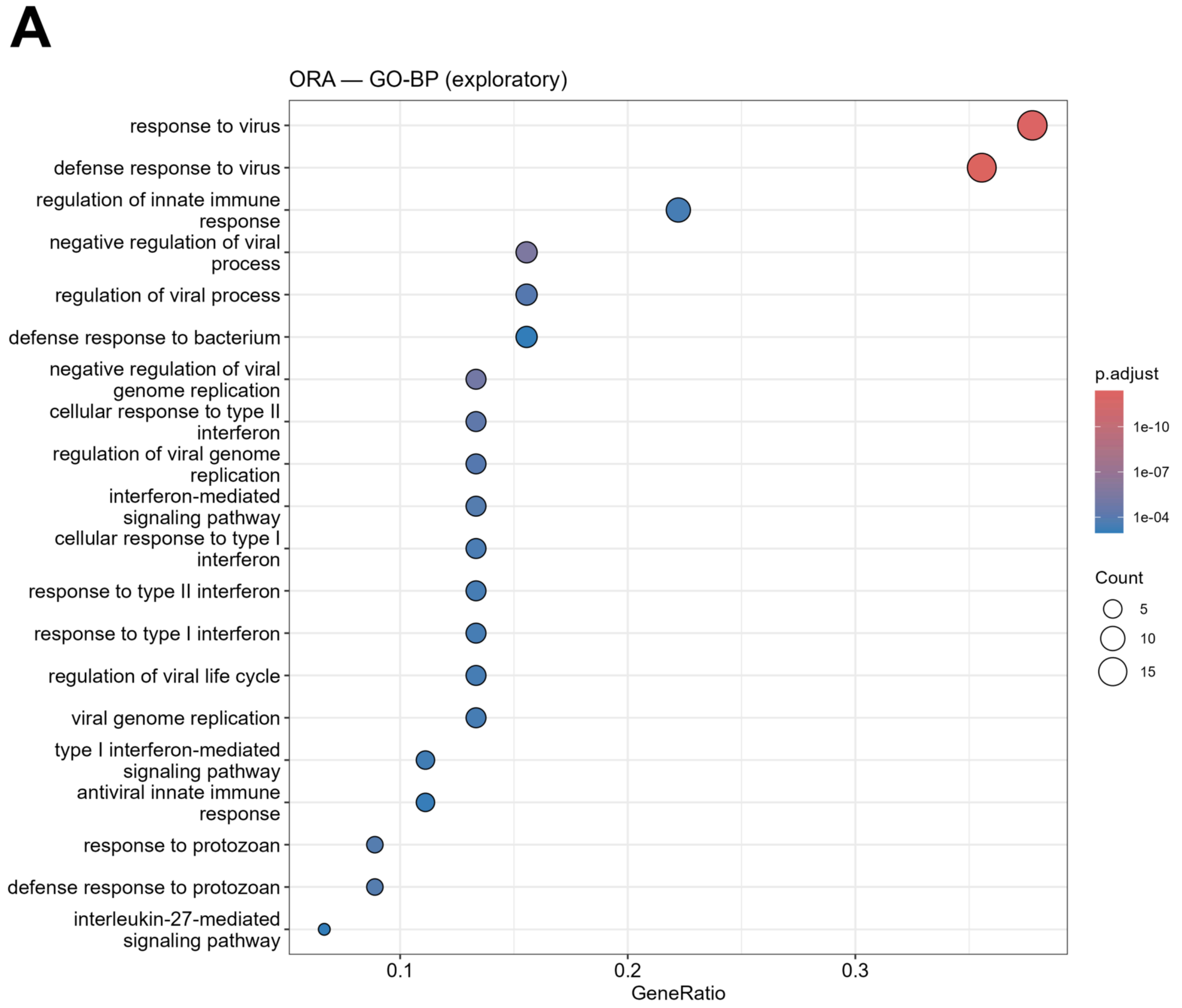


**Figure S9.** Full over-representation analysis of the 49 protein-coding consensus genes against GO Biological Process, before the simplification used in main Figure 4A.

#### Supplementary tables

Each table is a byte copy of a frozen analysis output; the Zenodo deposit records its source path, dimensions and SHA-256 checksum. The four tables affected by the v1.2 symbol correction, plus the v1.3 module annotation (S19) and GSE56153 alias comparison (S20), are reproduced in Appendix A; the remaining files follow the manuscript Data Availability statement.

| ID | Title | File(s) |
|---|---|---|
| **S1** | Core 50: per-study FDR and direction | S01a, S01b |
| **S2** | Broad consensus (932 genes) | S02 |
| **S3** | Core 50 and full meta-analysis | S03, S03b |
| **S4** | Per-gene external AUC, GSE28623 | S04 |
| **S5** | LASSO selection frequency | S05 |
| **S6** | PPI hub versus model-importance crosscheck | S06 |
| **S7** | Class-imbalance sensitivity | S07 |
| **S8** | MCPcounter differential abundance | S08 |
| **S9** | CIBERSORTx differential abundance | S09 |
| **S10** | MCPcounter–ssGSEA correlations | S10 |
| **S11** | Berry 393 platform mapping | S11 |
| **S12** | GSEA counts and GO concordance | S12a, S12b |
| **S13** | RF external validation and scale audit | S13a, S13b |
| **S14** | Berry direction reconstruction | S14a, S14b |
| **S15** | Secondary cohorts: scale and direction | S15a–S15c |
| **S16** | Discovery DE counts and selection facts | S16a, S16b |
| **S17** | Signature comparator, GSE28623 | S17 |
| **S18** | GSE107991 provenance | S18a, S18b |
| **S19** | Gene-level module annotation | S19 |
| **S20** | GSE56153 alias comparison | S20 |

#### Appendix A. Corrected and verification tables

Tables S04, S15b, S15c and S17 are the four v1.2 tables affected by the HGNC symbol correction. Tables S19 and S20 are v1.3 verification tables: S19 gives the gene-level module annotation rule and S20 shows the GSE56153 alias comparison. They are reproduced here so that the corrected values accompany the preprint. Values are rounded for display; the exact values are regenerated by the R12, R13, R18 and R19 scripts. The current v1.3.0 record contains the corrected tables; the earlier v1.0.0 version remains available and contains the earlier versions of the four v1.2 tables.

##### S04. Per-gene external AUC, GSE28623

Per-gene discriminative performance for the 50 core genes in GSE28623 (46 PTB, 37 HC), with DeLong confidence intervals and Benjamini–Hochberg-adjusted p values. Genes absent from the platform are marked NA. The corrected mapping recovers WARS1; three genes remain unrepresented.

| Gene | AUC | 95% CI low | 95% CI high | Raw p | Test | BH-adjusted p |
|---|---|---|---|---|---|---|
| OAS3 | 0.675 | 0.554 | 0.795 | 0.00456 | delong_z | 0.00565 |
| GBP6 | 0.565 | 0.441 | 0.690 | 0.304 | delong_z | 0.311 |
| CACNA1E | 0.468 | 0.340 | 0.596 | 0.621 | delong_z | 0.621 |
| IFI44L | 0.663 | 0.542 | 0.784 | 0.00817 | delong_z | 0.00985 |
| VAMP5 | 0.815 | 0.722 | 0.908 | 3.31e-11 | delong_z | 1.11e-10 |
| SLAMF8 | 0.731 | 0.620 | 0.842 | 4.44e-05 | delong_z | 6.96e-05 |
| TIMM10 | 0.763 | 0.654 | 0.872 | 2.11e-06 | delong_z | 3.97e-06 |
| OASL | 0.635 | 0.510 | 0.761 | 0.0342 | delong_z | 0.0392 |
| TCN2 | 0.752 | 0.646 | 0.858 | 3.45e-06 | delong_z | 6.01e-06 |
| SOCS1 | 0.687 | 0.571 | 0.804 | 0.00165 | delong_z | 0.00215 |
| ETV7 | 0.844 | 0.759 | 0.928 | 1.44e-15 | delong_z | 8.45e-15 |
| ASPHD2 | 0.818 | 0.727 | 0.910 | 7.57e-12 | delong_z | 2.96e-11 |
| PARP14 | 0.729 | 0.617 | 0.840 | 5.85e-05 | delong_z | 8.87e-05 |
| LAP3 | 0.790 | 0.690 | 0.889 | 1.25e-08 | delong_z | 3.46e-08 |
| SLC6A12 | 0.794 | 0.695 | 0.894 | 6.5e-09 | delong_z | 1.91e-08 |
| CLEC4D | 0.814 | 0.723 | 0.906 | 1.6e-11 | delong_z | 5.77e-11 |
| PSTPIP2 | 0.900 | 0.831 | 0.968 | 6.57e-30 | delong_z | 3.09e-28 |
| CMPK2 | NA | NA | NA | NA | not_in_validation | NA |
| P2RY14 | 0.784 | 0.683 | 0.884 | 3.13e-08 | delong_z | 7.74e-08 |
| SAMD9L | 0.818 | 0.729 | 0.908 | 3.75e-12 | delong_z | 1.6e-11 |
| XAF1 | 0.750 | 0.641 | 0.858 | 6.7e-06 | delong_z | 1.09e-05 |
| IFI35 | 0.702 | 0.583 | 0.822 | 0.000914 | delong_z | 0.00123 |
| MYL9 | 0.695 | 0.580 | 0.810 | 0.000889 | delong_z | 0.00123 |
| TRIM22 | 0.759 | 0.654 | 0.863 | 1.29e-06 | delong_z | 2.52e-06 |
| LHFPL2 | 0.883 | 0.805 | 0.961 | 7.39e-22 | delong_z | 1.16e-20 |
| IFI44 | 0.707 | 0.590 | 0.823 | 0.00049 | delong_z | 0.00071 |
| STAT1 | 0.783 | 0.683 | 0.884 | 2.82e-08 | delong_z | 7.36e-08 |
| PLSCR1 | 0.774 | 0.674 | 0.875 | 8.39e-08 | delong_z | 1.97e-07 |
| RTP4 | 0.751 | 0.643 | 0.858 | 4.56e-06 | delong_z | 7.66e-06 |
| WARS1 | 0.769 | 0.665 | 0.872 | 3.43e-07 | delong_z | 7.32e-07 |
| CEACAM1 | 0.848 | 0.765 | 0.932 | 2.12e-16 | delong_z | 1.43e-15 |
| HERC5 | 0.643 | 0.518 | 0.768 | 0.0255 | delong_z | 0.0299 |
| IFI27 | 0.575 | 0.446 | 0.705 | 0.252 | delong_z | 0.263 |
| FBXO6 | 0.839 | 0.753 | 0.925 | 1.05e-14 | delong_z | 5.5e-14 |
| ISG15 | 0.577 | 0.448 | 0.707 | 0.242 | delong_z | 0.258 |
| C1QB | 0.829 | 0.738 | 0.920 | 1.4e-12 | delong_z | 6.58e-12 |
| IFIT1 | 0.630 | 0.502 | 0.759 | 0.0465 | delong_z | 0.052 |
| CASP5 | 0.801 | 0.706 | 0.897 | 6.71e-10 | delong_z | 2.1e-09 |
| GBP4 | 0.757 | 0.654 | 0.861 | 1.12e-06 | delong_z | 2.29e-06 |
| GBP1 | 0.864 | 0.782 | 0.947 | 6.31e-18 | delong_z | 5.45e-17 |
| CXCL10 | 0.678 | 0.561 | 0.795 | 0.00281 | delong_z | 0.00357 |
| IDO1 | 0.583 | 0.453 | 0.712 | 0.21 | delong_z | 0.23 |
| RSAD2 | 0.711 | 0.592 | 0.830 | 0.000499 | delong_z | 0.00071 |

| Gene | AUC | 95% CI low | 95% CI high | Raw p | Test | BH-adjusted p |
|---|---|---|---|---|---|---|
| EPSTI1 | 0.776 | 0.671 | 0.880 | 2.32e-07 | delong_z | 5.19e-07 |
| IFIT3 | 0.761 | 0.652 | 0.871 | 3.13e-06 | delong_z | 5.66e-06 |
| FCGR1A | NA | NA | NA | NA | not_in_validation | NA |
| ANKRD22 | 0.861 | 0.779 | 0.944 | 6.95e-18 | delong_z | 5.45e-17 |
| GBP5 | 0.873 | 0.795 | 0.950 | 3.4e-21 | delong_z | 4e-20 |
| FCGR1CP | NA | NA | NA | NA | not_in_validation | NA |
| BATF2 | 0.876 | 0.801 | 0.951 | 7.85e-23 | delong_z | 1.84e-21 |

#### S15b. Secondary-cohort direction summary

Direction concordance in the two direction-only cohorts after the corrected symbol mapping. n_testable is the number of core genes measured; percent_concordant is the percentage whose mean difference matches the discovery direction.

| Dataset | PTB (n) | HC (n) | Testable genes | Concordant (%) |
|---|---|---|---|---|
| GSE34608 | 8 | 18 | 47 | 95.74 |
| GSE81746 | 4 | 2 | 48 | 95.83 |

#### S15c. Secondary-cohort per-gene direction

Per-gene mean expression in PTB and HC and the concordance flag for GSE34608 and GSE81746. discovery_direction is 1 for upregulated; concordant is TRUE when the sign of mean PTB minus mean HC matches it.

| Dataset | Gene | Discovery direction | Mean PTB | Mean HC | Concordant |
|---|---|---|---|---|---|
| GSE34608 | OAS3 | 1 | 4.160 | 3.143 | TRUE |
| GSE34608 | GBP6 | 1 | 7.376 | 7.319 | TRUE |
| GSE34608 | CACNA1E | 1 | 9.033 | 11.380 | FALSE |
| GSE34608 | IFI44L | 1 | 5.497 | 4.094 | TRUE |
| GSE34608 | VAMP5 | 1 | 5.630 | 5.133 | TRUE |
| GSE34608 | SLAMF8 | 1 | 3.712 | 3.236 | TRUE |
| GSE34608 | TIMM10 | 1 | 5.644 | 4.824 | TRUE |
| GSE34608 | OASL | 1 | 6.579 | 6.011 | TRUE |
| GSE34608 | TCN2 | 1 | 1.623 | 0.924 | TRUE |
| GSE34608 | SOCS1 | 1 | 2.537 | 3.039 | FALSE |
| GSE34608 | ETV7 | 1 | 3.253 | 2.038 | TRUE |
| GSE34608 | ASPHD2 | 1 | 2.500 | 2.263 | TRUE |
| GSE34608 | PARP14 | 1 | 4.252 | 3.295 | TRUE |
| GSE34608 | LAP3 | 1 | 7.265 | 5.267 | TRUE |
| GSE34608 | SLC6A12 | 1 | 2.111 | 1.478 | TRUE |
| GSE34608 | CLEC4D | 1 | 6.442 | 3.989 | TRUE |
| GSE34608 | PSTPIP2 | 1 | 2.231 | 1.210 | TRUE |
| GSE34608 | P2RY14 | 1 | 6.191 | 3.526 | TRUE |
| GSE34608 | SAMD9L | 1 | 6.564 | 4.946 | TRUE |
| GSE34608 | XAF1 | 1 | 6.010 | 4.844 | TRUE |
| GSE34608 | IFI35 | 1 | 4.842 | 4.700 | TRUE |
| GSE34608 | MYL9 | 1 | 2.164 | 2.123 | TRUE |
| GSE34608 | TRIM22 | 1 | 6.497 | 4.895 | TRUE |
| GSE34608 | LHFPL2 | 1 | 3.101 | 2.009 | TRUE |
| GSE34608 | IFI44 | 1 | 6.261 | 4.269 | TRUE |
| GSE34608 | STAT1 | 1 | 7.935 | 6.449 | TRUE |
| GSE34608 | PLSCR1 | 1 | 5.766 | 3.884 | TRUE |
| GSE34608 | RTP4 | 1 | 6.650 | 5.257 | TRUE |
| GSE34608 | WARS1 | 1 | 6.697 | 5.886 | TRUE |
| GSE34608 | CEACAM1 | 1 | 6.155 | 4.640 | TRUE |
| GSE34608 | HERC5 | 1 | 5.392 | 3.585 | TRUE |
| GSE34608 | IFI27 | 1 | 6.462 | 5.862 | TRUE |
| GSE34608 | FBXO6 | 1 | 3.719 | 3.057 | TRUE |
| GSE34608 | ISG15 | 1 | 8.801 | 7.794 | TRUE |
| GSE34608 | C1QB | 1 | 3.818 | 1.597 | TRUE |
| GSE34608 | IFIT1 | 1 | 8.472 | 6.330 | TRUE |
| GSE34608 | CASP5 | 1 | 7.710 | 6.678 | TRUE |
| GSE34608 | GBP4 | 1 | 6.652 | 5.771 | TRUE |
| GSE34608 | GBP1 | 1 | 6.325 | 3.989 | TRUE |
| GSE34608 | CXCL10 | 1 | 1.887 | 0.705 | TRUE |
| GSE34608 | IDO1 | 1 | 5.091 | 2.900 | TRUE |
| GSE34608 | RSAD2 | 1 | 5.400 | 2.888 | TRUE |
| GSE34608 | EPSTI1 | 1 | 5.529 | 2.781 | TRUE |

| Dataset | Gene | Discovery direction | Mean PTB | Mean HC | Concordant |
|---|---|---|---|---|---|
| GSE34608 | IFIT3 | 1 | 6.526 | 5.367 | TRUE |
| GSE34608 | ANKRD22 | 1 | 4.936 | 2.713 | TRUE |
| GSE34608 | GBP5 | 1 | 9.877 | 7.762 | TRUE |
| GSE34608 | BATF2 | 1 | 5.047 | 3.976 | TRUE |
| GSE81746 | OAS3 | 1 | 3.201 | 1.474 | TRUE |
| GSE81746 | GBP6 | 1 | 7.065 | 8.186 | FALSE |
| GSE81746 | CACNA1E | 1 | 4.030 | 1.788 | TRUE |
| GSE81746 | IFI44L | 1 | 5.601 | 1.887 | TRUE |
| GSE81746 | VAMP5 | 1 | 7.562 | 5.848 | TRUE |
| GSE81746 | SLAMF8 | 1 | 4.460 | 3.724 | TRUE |
| GSE81746 | TIMM10 | 1 | 5.372 | 4.842 | TRUE |
| GSE81746 | OASL | 1 | 7.092 | 5.940 | TRUE |
| GSE81746 | TCN2 | 1 | 2.183 | 1.119 | TRUE |
| GSE81746 | SOCS1 | 1 | 5.761 | 4.843 | TRUE |
| GSE81746 | ETV7 | 1 | 4.416 | 3.008 | TRUE |
| GSE81746 | ASPHD2 | 1 | 3.848 | 2.174 | TRUE |
| GSE81746 | PARP14 | 1 | 6.397 | 5.027 | TRUE |
| GSE81746 | LAP3 | 1 | 7.209 | 5.681 | TRUE |
| GSE81746 | SLC6A12 | 1 | 2.630 | 1.051 | TRUE |
| GSE81746 | CLEC4D | 1 | 6.513 | 4.548 | TRUE |
| GSE81746 | PSTPIP2 | 1 | 3.562 | 2.454 | TRUE |
| GSE81746 | CMPK2 | 1 | 6.493 | 5.513 | TRUE |
| GSE81746 | P2RY14 | 1 | 7.317 | 6.237 | TRUE |
| GSE81746 | SAMD9L | 1 | 8.065 | 6.786 | TRUE |
| GSE81746 | XAF1 | 1 | 5.334 | 3.789 | TRUE |
| GSE81746 | IFI35 | 1 | 5.958 | 5.114 | TRUE |
| GSE81746 | MYL9 | 1 | 3.983 | 2.997 | TRUE |
| GSE81746 | TRIM22 | 1 | 6.476 | 4.040 | TRUE |
| GSE81746 | LHFPL2 | 1 | 5.606 | 4.149 | TRUE |
| GSE81746 | IFI44 | 1 | 7.295 | 5.569 | TRUE |
| GSE81746 | STAT1 | 1 | 7.526 | 6.181 | TRUE |
| GSE81746 | PLSCR1 | 1 | 7.814 | 5.939 | TRUE |
| GSE81746 | RTP4 | 1 | 6.626 | 5.362 | TRUE |
| GSE81746 | WARS1 | 1 | 8.003 | 5.980 | TRUE |
| GSE81746 | CEACAM1 | 1 | 6.668 | 5.326 | TRUE |
| GSE81746 | HERC5 | 1 | 4.217 | 3.210 | TRUE |
| GSE81746 | IFI27 | 1 | 6.495 | 6.354 | TRUE |
| GSE81746 | FBXO6 | 1 | 5.062 | 3.772 | TRUE |
| GSE81746 | ISG15 | 1 | 9.710 | 8.553 | TRUE |
| GSE81746 | C1QB | 1 | 5.563 | 0.713 | TRUE |
| GSE81746 | IFIT1 | 1 | 8.695 | 7.886 | TRUE |
| GSE81746 | CASP5 | 1 | 9.338 | 8.402 | TRUE |
| GSE81746 | GBP4 | 1 | 7.511 | 5.803 | TRUE |
| GSE81746 | GBP1 | 1 | 6.193 | 4.193 | TRUE |
| GSE81746 | CXCL10 | 1 | 1.914 | 0.417 | TRUE |
| GSE81746 | IDO1 | 1 | 4.184 | 4.318 | FALSE |
| GSE81746 | RSAD2 | 1 | 5.499 | 3.839 | TRUE |
| GSE81746 | EPSTI1 | 1 | 4.592 | 2.633 | TRUE |
| GSE81746 | IFIT3 | 1 | 8.511 | 7.172 | TRUE |
| GSE81746 | ANKRD22 | 1 | 4.357 | 1.119 | TRUE |
| GSE81746 | GBP5 | 1 | 10.652 | 8.332 | TRUE |
| GSE81746 | BATF2 | 1 | 7.214 | 3.720 | TRUE |

#### S17. Signature comparator, GSE28623

Signature comparator on the same 83 GSE28623 samples. delong_p_vs_PTB50 is the paired DeLong test against the corrected 47-gene score; delong_p_vs_Sweeney and delong_p_vs_Berry_unsigned are the corresponding pairwise tests; delong_holm_primary3 is the Holm correction across the three primary contrasts against PTB_50. NA marks a test that is not defined for that row.

| Signature | Scoring | n | AUC | CI low | CI high | Wilcoxon p | DeLong p vs 50-gene | DeLong p vs Sweeney | DeLong p vs Berry unsigned | Holm p (primary 3) |
|---|---|---|---|---|---|---|---|---|---|---|
| PTB_50_ssGSEA | ssGSEA (47 genes) | 83 | 0.8302 | 0.7432 | 0.9172 | 5.912e-08 | NA | 0.01171 | NA | NA |
| Sweeney_3 | published directional score (log2) | 83 | 0.9125 | 0.8551 | 0.9698 | 1.202e-12 | 0.01171 | NA | 0.02715 | 0.02342 |
| Berry_unsigned_ssGSEA | ssGSEA | 83 | 0.8278 | 0.7396 | 0.9161 | 7.557e-08 | 0.8992 | 0.02715 | NA | 0.8992 |
| Berry_signed_ssGSEA_GSE19439 | signed ssGSEA (up ES - down ES), signs from GSE19439 | 83 | 0.9095 | 0.8443 | 0.9747 | 1.945e-12 | 0.002358 | 0.9011 | 0.00374 | 0.007074 |
| Berry_signed_ssGSEA_GSE19444 | signed ssGSEA (up ES - down ES), signs from GSE19444 | 83 | 0.9107 | 0.8462 | 0.9752 | 1.606e-12 | 0.001784 | 0.9407 | 0.003006 | NA |
| Berry_meanz_GSE19439 | mean z (up - down), signs from GSE19439 | 83 | 0.9136 | 0.8504 | 0.9768 | 9.894e-13 | 0.000979 | 0.9601 | 0.002275 | NA |
| Berry_meanz_GSE19444 | mean z (up - down), signs from GSE19444 | 83 | 0.9130 | 0.8493 | 0.9768 | 1.091e-12 | 0.001125 | 0.9803 | 0.00254 | NA |
| Four_gene_RF | frozen RF probability (log2 + z) | 83 | 0.8643 | 0.7851 | 0.9435 | 9.77e-10 | 0.1967 | 0.05574 | 0.2435 | NA |

#### S19. Gene-level module annotation

Module assignment for all 50 core genes under one documented rule. A gene is interferon if it belongs to the MSigDB Hallmark interferon-alpha or interferon-gamma union (v2023.2.Hs) or to the fixed five-gene ISG supplement (ETV7, GBP1, GBP5, HERC5, TRIM22); it is innate/myeloid if it belongs to the fixed eight-gene list; otherwise it is other. The table shows the Hallmark flags and the rule that fired for each gene.

| Gene | Module | Hallmark IFN-alpha | Hallmark IFN-gamma | Hallmark union | Curated ISG | Rule |
|---|---|---|---|---|---|---|
| ANKRD22 | Other | no | no | no | no | Other (no rule match) |
| ASPHD2 | Other | no | no | no | no | Other (no rule match) |
| BATF2 | Interferon | yes | yes | yes | no | Hallmark IFN-alpha/gamma union |
| C1QB | Innate / myeloid | no | no | no | no | Innate/myeloid list |
| CACNA1E | Other | no | no | no | no | Other (no rule match) |
| CASP5 | Innate / myeloid | no | no | no | no | Innate/myeloid list |
| CEACAM1 | Innate / myeloid | no | no | no | no | Innate/myeloid list |
| CLEC4D | Innate / myeloid | no | no | no | no | Innate/myeloid list |
| CMPK2 | Interferon | yes | yes | yes | no | Hallmark IFN-alpha/gamma union |
| CXCL10 | Interferon | yes | yes | yes | no | Hallmark IFN-alpha/gamma union |
| EPSTI1 | Interferon | yes | yes | yes | no | Hallmark IFN-alpha/gamma union |
| ETV7 | Interferon | no | no | no | yes | Curated ISG supplement |
| FBXO6 | Innate / myeloid | no | no | no | no | Innate/myeloid list |
| FCGR1A | Interferon | no | yes | yes | no | Hallmark IFN-alpha/gamma union |
| FCGR1CP | Innate / myeloid | no | no | no | no | Innate/myeloid list |
| GBP1 | Interferon | no | no | no | yes | Curated ISG supplement |
| GBP4 | Interferon | yes | yes | yes | no | Hallmark IFN-alpha/gamma union |
| GBP5 | Interferon | no | no | no | yes | Curated ISG supplement |
| GBP6 | Interferon | no | yes | yes | no | Hallmark IFN-alpha/gamma union |
| HERC5 | Interferon | no | no | no | yes | Curated ISG supplement |
| IDO1 | Interferon | no | yes | yes | no | Hallmark IFN-alpha/gamma union |
| IFI27 | Interferon | yes | yes | yes | no | Hallmark IFN-alpha/gamma union |
| IFI35 | Interferon | yes | yes | yes | no | Hallmark IFN-alpha/gamma union |
| IFI44 | Interferon | yes | yes | yes | no | Hallmark IFN-alpha/gamma union |
| IFI44L | Interferon | yes | yes | yes | no | Hallmark IFN-alpha/gamma union |
| IFIT1 | Interferon | no | yes | yes | no | Hallmark IFN-alpha/gamma union |
| IFIT3 | Interferon | yes | yes | yes | no | Hallmark IFN-alpha/gamma union |
| ISG15 | Interferon | yes | yes | yes | no | Hallmark IFN-alpha/gamma union |
| LAP3 | Interferon | yes | yes | yes | no | Hallmark IFN-alpha/gamma union |
| LHFPL2 | Other | no | no | no | no | Other (no rule match) |
| MYL9 | Other | no | no | no | no | Other (no rule match) |
| OAS3 | Interferon | no | yes | yes | no | Hallmark IFN-alpha/gamma union |
| OASL | Interferon | yes | yes | yes | no | Hallmark IFN-alpha/gamma union |
| P2RY14 | Interferon | no | yes | yes | no | Hallmark IFN-alpha/gamma union |
| PARP14 | Interferon | yes | yes | yes | no | Hallmark IFN-alpha/gamma union |
| PLSCR1 | Interferon | yes | yes | yes | no | Hallmark IFN-alpha/gamma union |
| PSTPIP2 | Innate / myeloid | no | no | no | no | Innate/myeloid list |
| RSAD2 | Interferon | yes | yes | yes | no | Hallmark IFN-alpha/gamma union |
| RTP4 | Interferon | yes | yes | yes | no | Hallmark IFN-alpha/gamma union |
| SAMD9L | Interferon | yes | yes | yes | no | Hallmark IFN-alpha/gamma union |
| SLAMF8 | Innate / myeloid | no | no | no | no | Innate/myeloid list |
| SLC6A12 | Other | no | no | no | no | Other (no rule match) |
| SOCS1 | Interferon | no | yes | yes | no | Hallmark IFN-alpha/gamma union |
| STAT1 | Interferon | no | yes | yes | no | Hallmark IFN-alpha/gamma union |
| TCN2 | Other | no | no | no | no | Other (no rule match) |
| TIMM10 | Other | no | no | no | no | Other (no rule match) |
| TRIM22 | Interferon | no | no | no | yes | Curated ISG supplement |
| VAMP5 | Interferon | no | yes | yes | no | Hallmark IFN-alpha/gamma union |
| WARS1 | Interferon | yes | yes | yes | no | Hallmark IFN-alpha/gamma union |
| XAF1 | Interferon | no | yes | yes | no | Hallmark IFN-alpha/gamma union |

#### S20. GSE56153 alias comparison

ssGSEA performance in GSE56153 before and after restoring the legacy symbols WARS1 and IDO1. The alias-corrected run used 47 genes; the old-symbol run used 45. AUC, DeLong confidence interval and Wilcoxon p value are identical; the correction shifted 10 of 36 individual ssGSEA ranks by at most two positions (Spearman rho = 0.9979) without changing the case-control discrimination. Values are rounded for display.

| Dataset | Version | Genes | Absent | PTB (n) | HC (n) | AUC | CI low | CI high | Wilcoxon p |
|---|---|---|---|---|---|---|---|---|---|
| GSE56153 | old_symbols | 45 | ETV7;CMPK2;WARS1;IDO1;FCGR1CP | 18 | 18 | 0.9414 | 0.8677 | 1.0000 | 4.59e-07 |
| GSE56153 | alias_corrected | 47 | ETV7;CMPK2;FCGR1CP | 18 | 18 | 0.9414 | 0.8677 | 1.0000 | 4.59e-07 |
